# Leveraging learned representations and multitask learning for lysine methylation site discovery

**DOI:** 10.1101/2025.08.27.672583

**Authors:** François Charih, Mullen Boulter, Kyle K. Biggar, James R. Green

## Abstract

Lysine methylation is a dynamic and reversible post-translational modification of proteins carried out by lysine methyltransferase enzymes. The role of this modification in epigenetics and gene regulation is relatively well understood, but our understanding of the extent and the role of lysine methylation of non-histone substrates remains fairly limited. Several lysine methyltransferases which methylate non-histone substrates are overexpressed in a number of cancers and are believed to be key drivers of cancer progression. There is great incentive to identify the lysine methylome, as this is a key step in identifying drug targets. While numerous computational models have been developed in the last decade to identify novel lysine methylation sites, the accuracy of these model has been modest, leaving much room for improvement. In this work, we leverage the most recent advancements in deep learning and present a transformer-based model for lysine methylation site prediction which achieves state-of-the-art accuracy. In addition, we show that other post-translational modifications of lysine are informative and that multitask learning is an effective way to integrate this prior knowledge into our lysine methylation site predictor, MethylSight 2.0. Finally, we validate our model by means of mass spectrometry experiments and identify 68 novel lysine methylation sites. This work constitutes another contribution towards the completion of a comprehensive map of the lysine methylome.

## Introduction

Lysine methylation extends far beyond the realm of histone proteins and that it may be more prevalent than previously believed (Biggar and Li 2015). Studies have uncovered the involvement of non-histone lysine methylation in oncogenic processes chemoresistance and cancer cell proliferation (Carlson and Gozani 2016; Han et al. 2019; Huang et al. 2024), making it a very attractive target for anti-cancer therapies. Therapies targeting non-histone KMTs are emerging, with some even having reached the clinical trial stage with promising signs of efficacy (Feoli et al. 2022; Straining and Eighmy 2022). For instance, Tazemetostat, an inhibitor of EZH2, was trialed and received approval for the treatment of blood and solid malignancies (Straining and Eighmy 2022). EZH2 promotes tumorigenesis in glioblastoma and prostate cancer models via STAT3 methylation (Kim et al. 2013; Xu et al. 2012) and in diffuse large B-cell and follicular lymphomas via methylation of the PRC2 complex (Straining and Eighmy 2022). Considering that many cancers are driven by KMT overexpression, uncovering the human lysine methylome and the associated KMTs/KDMs would have profound implications in drug discovery and facilitate the identification of drug targets for therapeutic intervention, but would also broaden our general understanding of the lysine methylation-dependent biological processes.

Identification of novel lysine methylation sites is possible through experimental methods such as mass spectrometry (MS) (Lanouette et al. 2014). That said, the process is sufficiently resource-intensive that identifying new sites at the proteome scale experimentally is logistically impractical. For that reason, a number of machine learning prediction models have been developed over the years to tackle the lysine methylation prediction problem. The rationale behind these models is that computation can guide our efforts so that time and resources can be invested on validating the most promising potential lysine methylation sites.

A wide range of machine learning models trained on lysine methylation datasets to predict lysine methylation sites from sequence only have been published over the last two decades. Most of them have relied on the use of “traditional” machine learning algorithms such as support vector machines (SVMs) or random forests (RFs) and human-crafted numerical features. The first predictor, MeMo (Chen et al. 2006), was built using an SVM classifier using what is now refered to as “one-hot” encoding for a 15 amino acids lysine-centered window. The training set used in that study consisted of a total of 156 positive lysine methylation sites, which represents only an infinitesimal fraction of the space of all lysine-centered 15-mers (156/14^20^ ≈ 10^−19^% of the space of possible windows). More recent models used different strategies to represent lysine-centered windows. For example, iMethyl-PseAAC (Qiu et al. 2014) used a SVM model in conjunction with a representation the authors termed “pseudo amino acid composition” (PseAAC). This representation combines evolutionary information from the position-specific scoring matrix (PSSM), physicochemical properties of individual amino acids from the AAIndex (Kawashima et al. 2008), and disorder scores to generate a 346-dimensional feature vector. Another well-cited method is GPS-MSP (Group-based Prediction System Methyl-group Specific Predictor), an algorithm published in 2017 (Deng et al. 2017), was trained on 1,521 methyllysines sites and used an alignment-based custom scoring function to measure window similarity in conjunction with *k*-means clustering (unsupervised learning) to predict not only methylation sites, but also the methylation state (mono-, di-, or tri-), an ambitious task given the scarcity of data available to train models to this level of granularity.

Attempts to leverage deep learning methods to address the challenging task of accurately predicting the lysine methylome remain scarce as of the time of writing. Recently, Spadaro *et al*. applied convolutional neural networks (CNNs) to representations combining phylogenetic, physicochemical, structural, and binary encodings to predict lysine methylation sites (Spadaro, Sharma, and Dehzangi 2024). The PTM-Mamba model (Peng et al. 2025) makes use of the Mamba architecture, an attention-free state-space model with architectural optimizations designed to enhance computational efficiency over long sequences. More specifically, PTM-Mamba fuses Mamba-generated sequence embeddings with embeddings generated by the ESM-2 protein language model (pLM) to predict a wide array of PTMs which, for some reason, do not include lysine methylation.

Bepler and Berger have shown that multitask language models better capture the semantic organization of proteins by training a bidirectional long short-term memory (LSTM) to complete three tasks simultaneously: masked language modeling, residue-residue contact prediction, and structural similarity prediction (Bepler and Berger 2021).

Here, we hypothesize that knowledge about other PTMs of lysine residues such as acetylation, ubiquitination, sumoylation, and phosphorylation can inform models designed to predict methylation. This hypothesis arises as a result of the fact that enzymes catalyzing PTMs are known to compete to modify the same lysine residues and that this competition constitutes an additional layer of protein function modulation. Consistent with this fact, the PhosphoSitePlus (Hornbeck et al. 2015), a well-maintained database which curates post-translational modifications of proteins, lists 31,845 lysine residues in human proteins with annotations for two or more post-translational modifications (PTMs) (Figure 1). Of the 4,958 validated methylation sites in the database, 2,375 sites (48%) are subject to at least one other known modification.

**Figure 1.**
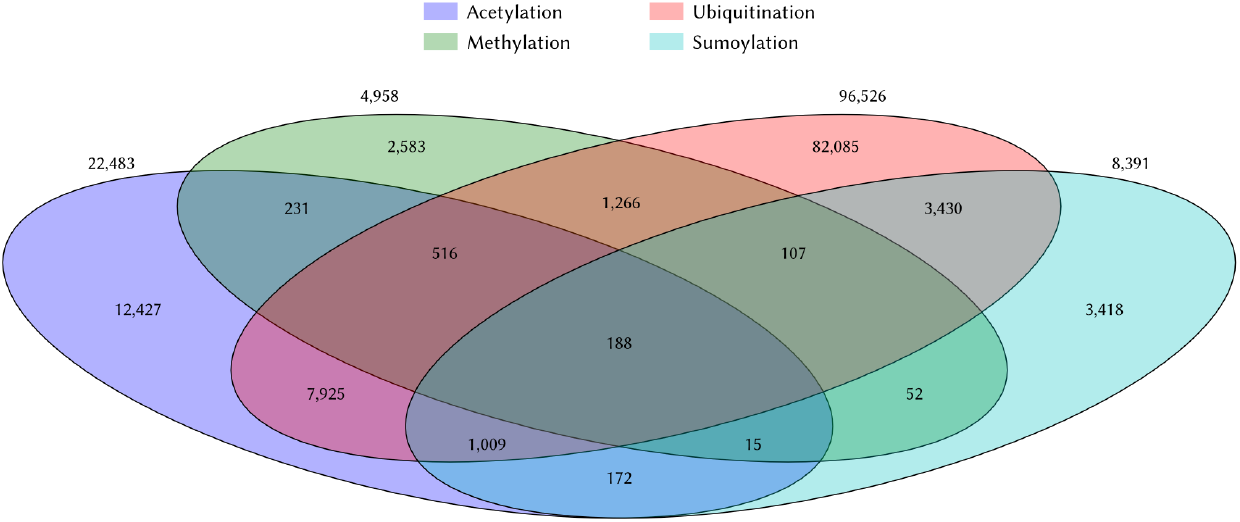
Co-occurence of common post-translational modification of lysines in the Phospho-SitePlus database. The overlap of four major PTMs of lysines among human proteins are shown in this Venn diagram. These numbers were computed using the 10/17/24 update of the PhosphoSitePlus database (Hornbeck et al. 2015). Many yet-to-be-discovered modifications remain to be deposited. Of the four major modifications of lysines, methylation is the one with the fewest annotations.

**Figure 2.**
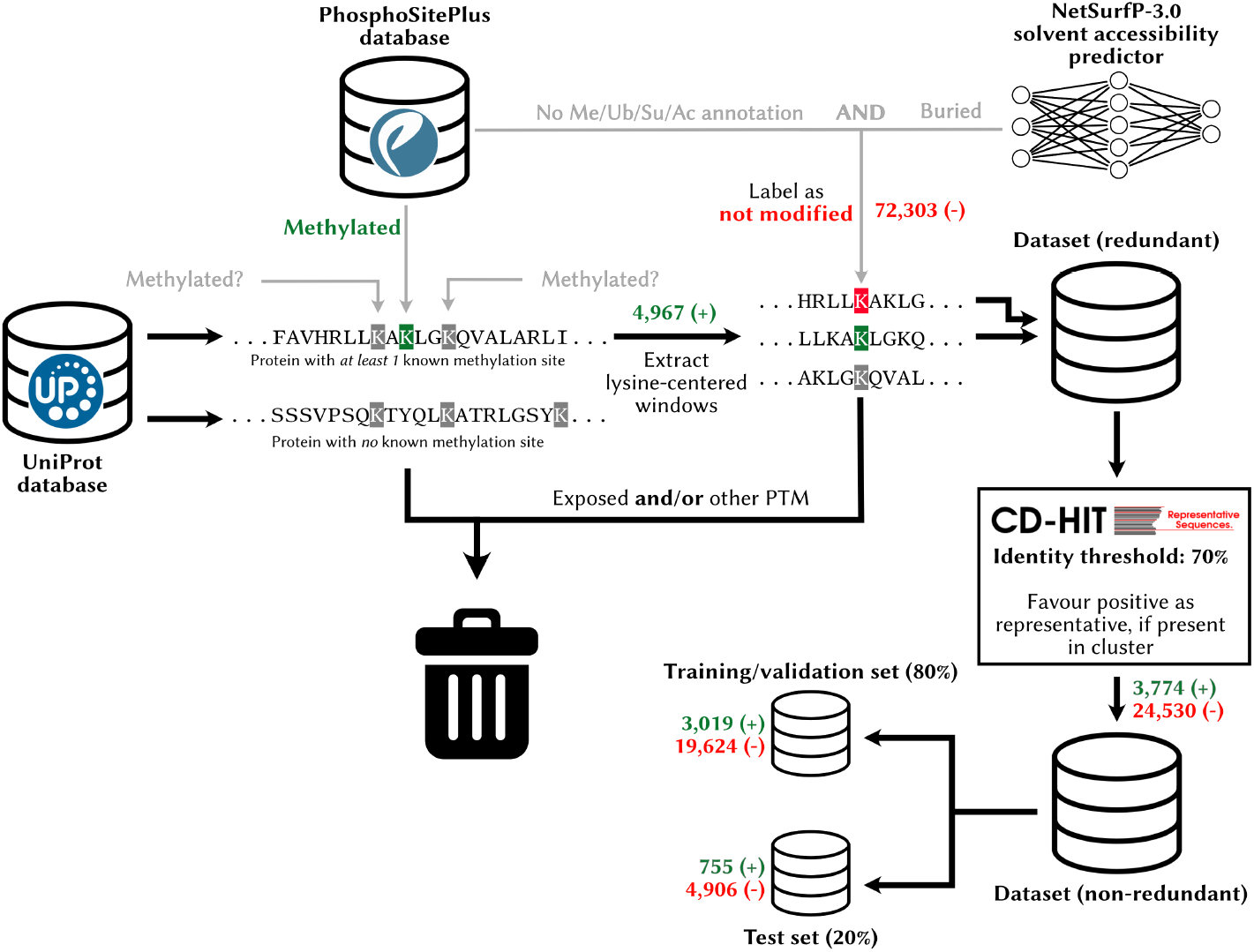
Preparation of a high-quality lysine modification dataset. To create the dataset used as part of this study, we sourced data from the PhosphoSitePlus database (for PTM annotations) and the UniProt database (for protein sequences). Only proteins with at least one methylated lysine residue were included in the dataset. Exposed residues of unknown status and/or having an anotation for another PTM were discarded, while the remaining lysines not known to be methylated were selected to make up the negative training data. The redundancy in the dataset was reduced with CD-HIT, using a window size of 31 for clustering and a 70% identity threshold. A blind test set was created by setting aside 20% of this data.

It is a reasonable supposition that there might be some partial overlap in the physico-chemical environments that promote lysine methylation and other PTMs, such as solvent accessibility, surrounding amino acid properties, and steric (spatial) constraints. As a result, the lysine methylation prediction problem is amenable to a multitask learning formulation, wherein lysine methylation prediction is one task among several PTM prediction tasks, and that jointly training a single model on several such tasks concomitantly could lead to better prediction accuracy through knowledge transfer.

To our knowledge, this idea has been exploited only once for the uncommon *propionylation* PTM of lysines (Li et al. 2021). In that work, the authors trained a recurrent neural network (RNN) on lysine malonylation sites and *fine-tuned* it using a dataset of lysine propionylation sites to extract features that are then fed into an SVM classifier. That work did exploit transfer learning, but did not use a multitask learning scheme, as the training was not performed for both tasks (*i*.*e*. propionylation and malonylation prediction) *simultaneously*. A model was trained for the malonylation prediction task first, and subsequently trained to predict propionylation sites, of which there were fewer known instances (431 as opposed to 9,584).

Currently, no one has proposed a lysine methylation site prediction model that leverages 1) state-of-the-art neural architecture, *i*.*e*. the transformer *and* 2) domain adaptation by means of transfer learning techniques such as multitask learning. In addition, very few groups have proven with *in vitro* experiments that the estimates of accuracy of their models translate into the lab upon deployment.

In this work, we address these opportunities to develop a more accurate and robust lysine methylation predictor. Our contributions are summarized below.

#### Contribution 1 - Improved prediction accuracy

We bootstrap embeddings generated with pLMs trained on millions of protein sequences to train a model, MethylSight 2.0, which produces *dramatically* more accurate predictions than previous lysine methylation predictors;

#### Contribution 2 - Use of multitask learning to enhance model accuracy

We demonstrate that training a transformer-based neural network architecture with a multitask learning strategy can lead to more accurate predictions;

#### Contribution 3 - Experimental validation

We show, by means of MS validation experiments performed on sites predicted to be methylated by our model, MethylSight 2.0, that the accuracy our model translates experimentally and identify 68 novel lysine methylation sites;

#### Contribution 4 - Bioinformatics analysis of the MethylSight 2.0 predicted lysine methylome

We deploy MethylSight 2.0 at the proteome scale to identify previously unknown methylation sites and conduct analyses to identify biological processes wherein lysine methylation sites are overrepresented.

## Methods

### Lysine methylation dataset preparation

With the intent of creating a dataset for multitask learning involving multiple PTM types, we retrieved PTM data for lysines occurring in human proteins by mining the PhosphoSitePlus database (Hornbeck et al. 2015) (10/17/24 update) for methylation, ubiquitination, sumoylation, and acetylation, which are all known to be modifications of lysines. The composition of the PhosphoSitePlus dataset is summarized in Table 1.

**Table 1.** Composition of the PhosphoSitePlus dataset (human lysines)

| Modification | Important functions | Unique proteins | Positive sites | Negative sites (low confidence) |
| --- | --- | --- | --- | --- |
| Methylation | Chromatin and gene expression regulation (histones), signaling, enzyme (in)activation (Lukinović et al. 2020) | 2,751 | 4,966 | 157,385 |
| Acetylation | Protein stability, regulation of PPIs and protein-DNA interactions (histones) (Narita, Weinert, and Choudhary 2019) | 7,047 | 22,547 | 333,013 |
| Sumoylation | Alteration of molecular interactions of substrate through addition/hiding interaction surfaces (Geiss-Friedlander and Melchior 2007) | 2,646 | 8,391 | 124,274 |
| Ubiquitination | Regulation of protein degradation, autophagy, protein trafficking (Damgaard 2021) | 11,712 | 96,545 | 377,547 |

Gathering positive lysine modification data is relatively straightforward, but identifying the “negative” sites to complete the training set needed to train a binary classifier is more arduous. It is difficult to ascertain with confidence that a lysine not known to be modified *never* is. In reality, sites taken to be “negative” may correspond to yet-to-be discovered methylation sites. Some groups simply take as negative training examples sites not known to be modified (Shrestha et al. 2024; Zheng et al. 2020), which we argue might disproportionately bias the learning algorithm towards making negative predictions. For this reason, it is typical to only use a subset of sites without PTM annotations as negative in PTM site prediction challenges, following some heuristics (Biggar et al. 2020). A typical practice is to only label as negatives unlabeled sites that occur within a protein containing a known modified site elsewhere (Biggar et al. 2020; Chen et al. 2006; Shi et al. 2012; Shi et al. 2015; Xue et al. 2005). To train and test MethylSight (Biggar et al. 2020), a lab-validated SVM-based model that achieved state-of-the-art performance upon publication, we applied two additional criteria to label potential lysine methylation as “negative” in addition to the latter. More specifically, sites were considered “negative” in the training set if they were not known to be substrates for another PTM (ubiquitination, sumoylation, or acetylation) *and* were predicted to be buried, *i*.*e*. to have a relative solvent accessibility factor < 0.2, as predicted with NetSurfP v1.0 (Petersen et al. 2009). In this work, we applied the same curation method, but used NetSurfP v3.0 (Høie et al. 2022). This approach allowed us to build a dataset with high-confidence negatives.

To address the issue of redundancy in the data, which could cause overrepresentation of certain patterns in the dataset and data leakage, *i*.*e*. similar patterns in the training and test data, we clustered the windows based on sequence identity at a similarity threshold of 70% with CD-HIT (Fu et al. 2012), as done previously (Biggar et al. 2020), and selected one representative from each cluster at random, favouring a positive representative (methylation site) if one occurred within a cluster. Finally, 20% of the non-redundant sites were set aside for testing.

We used an identical workflow to assemble the training sets for ubiquitination, acetylation, and sumoylation sites required for multitask learning, but did not set any data aside for testing, given that we are only interested in methylation site prediction.

The composition of the final dataset is presented in Table 2.

**Table 2.** Composition of the high-confidence dataset used to train and test the models.

|  | Methylation | Ubiquitination | Acetylation | Sumoylation |
| --- | --- | --- | --- | --- |
| Training set (pos/neg) | 2,415/15,699 | 68,539/58,314 | 15,791/51,498 | 5,737/15,862 |
| Validation set (pos/neg) | 604/3,925 |  |  |  |
| Test set (pos/neg) | 755/4,906 |  |  |  |

### Pre-trained protein-language model embeddings

pLMs have been shown to generate rich embeddings that capture physicochemical, phylogenetic and structural information that are extremely useful for a variety of downstream tasks, including structure prediction (Avraham et al. 2023; Lin et al. 2023), property prediction (*e*.*g*. viscosity (Hao and Fan 2024), stability (Schmirler, Heinzinger, and Rost 2024), *etc*.), localization prediction (Elnaggar et al. 2023; Schmirler, Heinzinger, and Rost 2024), and peptide binder design (Bhat et al. 2025; Brixi et al. 2023; Chen et al. 2025), to cite a few.

Given that these representations were learned on massive collections of protein sequences and performed well on these tasks, we hypothesized that they may also contain useful information for the prediction of lysine methylation sites. Moreover, these embeddings capture more context about the potential methylation sites than traditional human-engineered representations. They consider a large portion (or all) of the protein, depending on the pLM’s context length (window size) and the protein length.

We leveraged representations learned by three state-of-the-art foundational model pLMs: ProtT5 (Elnaggar et al. 2021), ESM-2 (Lin et al. 2023), and Ankh (Elnaggar et al. 2023) (Table 3) and fed the human proteome to all three models to generate embeddings for each lysine in the training and test sets – and to later predict the comprehensive human lysine methylome.

**Table 3.** Foundational protein language models used to embed potential lysine methylation sites.

| Model | Architecture | Version | Embedding dimension | Parameters (approx.) | Training strategy | Training data |
| --- | --- | --- | --- | --- | --- | --- |
| ProtT5 (Elnaggar et al. 2021) | Encoder-decoder | ProtT5-XL-BFD | 1,024 | 3B | 1-gram random masking with demasking | BFD (pre-training; ~2.1B sequences) and UniRef50 (fine-tuning; ~45M sequences) |
| ESM-2 (Lin et al. 2023) | Encoder-only | ESM-2-T33-650M-UR50D | 1,280 | 650M | 1-gram random masking with demasking | UniRef50+90 (~65M sequences) |
| Ankh (Elnaggar et al. 2023) | Encoder-decoder | Ankh Large | 1,536 | 1B | 1-gram random masking with full sequence reconstruction | UniRef50 (~45M sequences) |

### Training multilayer perceptrons leveraging pLM-generated embeddings

Using Optuna (Akiba et al. 2019), we trained 50 MLPs (MLPs) on the embedding vectors of lysine sites extracted with ProtT5, ESM-2 and Ankh, sampling at random the learning rate, number and width of of hidden layers, and the dropout rate used for regularization. We selected as our final models the ones with the highest validation area under the precision-recall curve (AUPRC). All models were trained using PyTorch (Paszke et al. 2019) with the Adam optimizer (Kingma and Ba 2017) with a batch size of 64, using binary cross-entropy as the loss function:

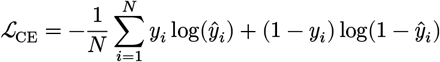

where *y*_*i*_ = 1 if the site *i* is methylated and *y*_*i*_ = 0 otherwise, while *ŷ*_*i*_ ∈ [0, 1] is the predicted probability of that the site *i* is methylated.

We selected the model using an early stopping strategy, using the validation loss to monitor for overfitting. We repeated this procedure using the embeddings generated by all three aforementioned pLMs. In addition, we trained MLPs on a “combined” representation resulting from a concatenation of all three embeddings, for a total of four final MLP models.

### Training a transformer model

To determine whether training a transformer-based model could further improve the quality of the predictions, we implemented a model which leverages this architecture.

We used a context size of 31 amino acids, the ProtT5 embeddings as representations for the individual amino acids in the sequence (*i*.*e*. the tokens), and padded with a zero-filled 1,024-D vector, if the lysine site was too close to the end of the protein chain (Figure 3A). To capture the positional information of the individual tokens, we used the cannonical positional embedding strategy described in (Vaswani et al. 2017). We used 4 heads in each attention block. A schematic representation of this architecture is shown in Figure 3B. The Adam optimizer was used, but with a batch size of 128.

**Figure 3.**
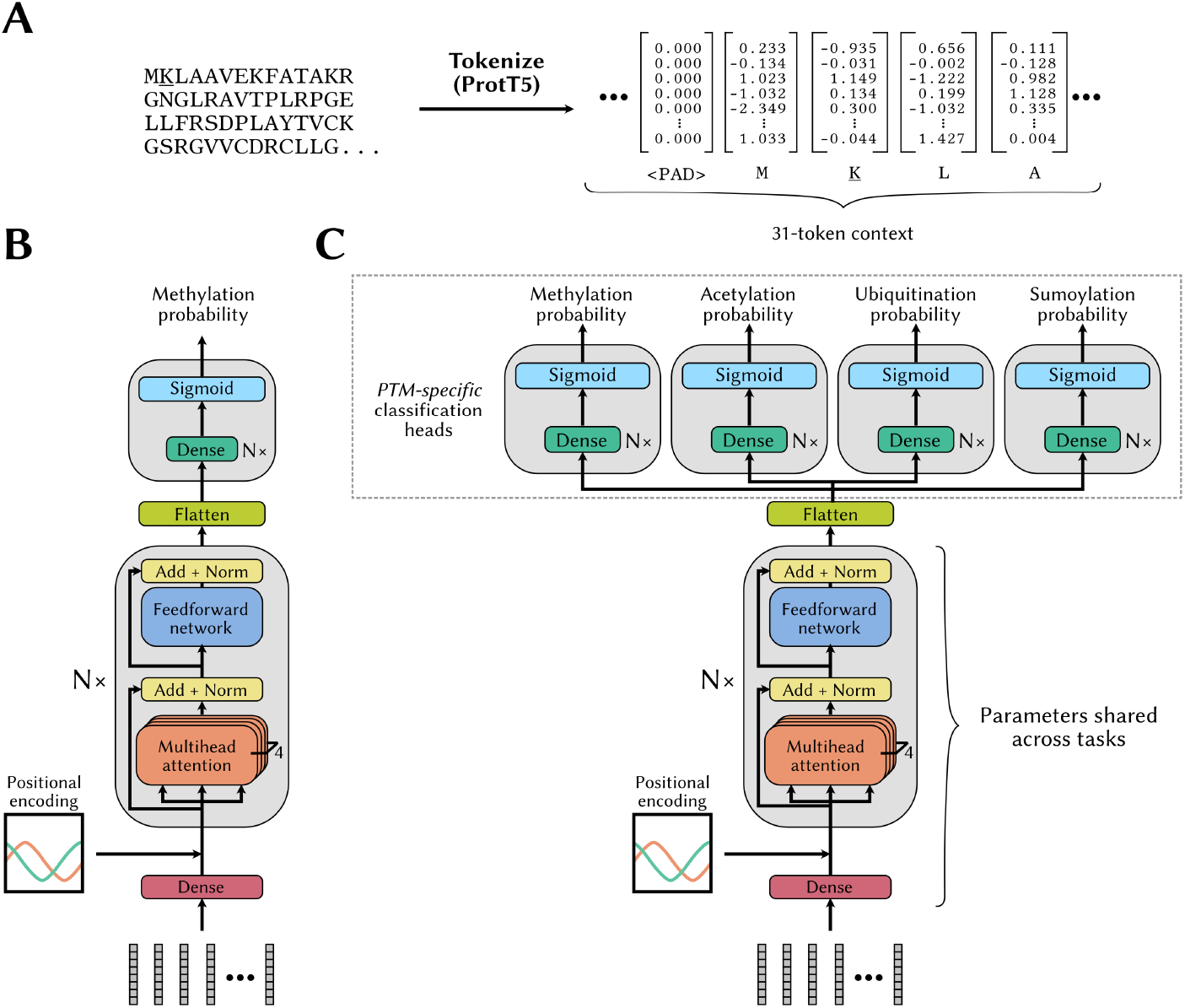
Transformer model architectures for methylation site prediction without and with multitask learning. **(A)** The inputs to our transformer-based models are the ProtT5 embeddings extracted from the full protein, with a context window of 31, centered around the lysine residue of interest. When a lysine residue is too close to an end of the protein sequence, null embeddings (<PAD>) are appended to complete the context window. **(B)** Architecture of our vanilla transformer-based model for methylation probability prediction. **(C)** Modified transformer architecture designed to enable a multitask learning strategy. More specifically, after the flattening layer, instance representations are sent to one of four PTM-specific classification heads, depending on the task associated with the individual instances.

Similarly to the approach used to train the MLPs, we conducted hyperparameter tuning in a randomized fashion and varied the number of encoder transformer blocks, the learning rate, the number of hidden layers in the classification module (*i*.*e*. the dense layers that follow the transformer layers), and the width of the “embedding layer” and trained a total of 50 models.

We used the same loss function and early stopping stratedy as for the MLPs to select the final model for each run. The final transformer architecture selected was the one with the highest AUPRC on the validation set.

### Multitask learning with a transformer model

To investigate whether a multitask learning strategy could enhance the quality of the predictions, we enriched our training set with sites and their annotation for the three other PTMs of interest: acetylation, ubiquitination, and sumoylation.

In this context, the “tasks” consist in predicting the four different PTMs of interest. We do not know or can’t assume with a satisfying level of certainty the true label for each task for all sites. For example, we may know that a site is acetylated, but not know whether it is also ubiquitinated. Consequently, we chose to not associate each instance or site with 4 labels. Instead, each instance in the dataset is a site associated with a PTM and a label associated to that site and PTM. Consequently, a given site may appear up to 4 times in the training set, in the specific case where a label for each PTM is known.

We implemented another transformer model where the last transformer block is followed by a flattening layer whose output is sent to one of four classification heads, depending on the task, *i*.*e*. prediction of methylation, acetylation, *etc*. (Figure 3C). Each classification head is designed to predict whether a site is subjected to the corresponding PTM. For each instance in the training set, we only probe the probability output by the head corresponding to the PTM (“task”) associated with the instance.

We use a custom batch sampling strategy to train the model wherein all instances in a batch are associated with only *one* of the four PTM. This ensures that the loss over a batch is only used to update the parameters of the classification head associated with the PTM (and the upstream parameters), but not the three classification heads which are used for the other tasks. In other terms, we use *partial* parameter sharing, *i*.*e*. only the parameters in the transformer layers and upstream are shared across tasks.

Given that the methylation sites are vastly outnumbered by other sites, we multiply the loss for methylation batches by a factor *γ* in order to produce larger updates for the shared model weights when methylation sites are misclassified relative to misclassified instances of other PTMs. We tried *γ* ∈ {1, 13.5, 20}, 1:13.5 being approximatively the methylation-to-other PTMs ratio.

The loss function for the multitask learning strategy effectively takes the form:

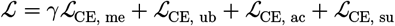

where

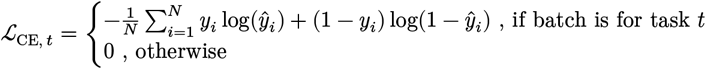

The rest of the model selection was done as for the transformer model without the multitask learning training strategy described in the previous section. We henceforth refer to this model, *i*.*e*. the transformer-style architecture trained with a multitask learning strategy on ProtT5 embeddings, as **MethylSight 2.0**.

### Estimation of the expected imbalance

To accurately estimate the precision of MethylSight 2.0 upon deployment on the human proteome, an estimate of the class imbalance is required. The human proteome in UniProt/Swiss-Prot database (2025_01 release) (The UniProt Consortium 2025) comprises 654,185 lysines residues in 20,417 unique proteins, of which an unknown fraction can be methylated under specific biological circumstances such as in response to a biological event, in a stage of development, or in specific tissue types.

Berryhill *et al*. (Berryhill et al. 2023) published a study which provides some useful insight into the ratio of methylated to unmethylated lysines observable through mass spectrometry experiments. In their study, they assessed the sequence bias of commercially available pan-methyllysine antibodies and performed global profiling of lysine methylation in HEK293T (human embryonic kidney) and U2OS (human osteosarcoma) cells with samples enriched with combinations of less biased anti-Kme1, anti-Kme2, and anti-Kme3 antibodies and their combinations. They identified a total of 5,089 lysine methylation sites evenly distributed through the proteome, of which 4,862 are novel.

Using the data collected in this study, we made the assumption that the estimated the imbalance ratio of methylated-to-unmethylated lysines *detectable via mass spectrometry without and following enrichment* with the antibodies currently in use to be roughly 1:36. This corresponds to the ratio of methylated lysines to lysines not found to be methylated in the proteins that were pulled down in the samples (*i*.*e*. with at least one epitope for the anti-Kme antibodies used). It is difficult to speculate about what lysines are or are not methylated in proteins that were not pulled down, so we only estimate what one may observe in a global profiling experiment with mass spectrometry. We use this ratio to evaluate the anticipated precision of MethylSight-2.0, when coupled with a mass spectrometry experiment.

This figure is an approximation derived from samples extracted from two specific cell types, and as such, it may not apply uniformly across all tissue types and across the entire proteome.

### Selection of predicted methylation sites for *in vitro* validation

We subsequently sought to estimate the actual precision of MethylSight 2.0 upon deployment onto the human proteome. To achieve this, we selected 100 sites predicted to be methylated by MethylSight 2.0, but which were not known methylation sites. Using a conservative threshold on predicted methylation probability (*i*.*e*. PCPr_1:36_ = 0.75), we sampled 50 sites at random from each of the following two sets:

1. **Set 1:** Exposed lysine residues known to be acetylated, ubiquitinated, and/or sumoylated;
2. **Set 2:** Exposed lysine residues with no known modification.

Furthermore, under the hypothesis supported by the phenomena of PTM competition that these other modifications provide useful information for the identification of novel lysine methylation sites, one would expect to detect more methylation events within sites sampled from Set 1 than within sites sampled from Set 2. To allow for this comparison, we ensured that the methylation probabilities were similarly distributed in both samples.

### Validation of predicted methylation sites via mass spectrometry

Using the Pyteomics package for Python (Levitsky et al. 2018), we generated an isolation list tabulating the peptide fragments and their mass-to-charge ratios for the +2, +3 and +4 charged states and for all four methylation states (null, mono-, di-, tri-), resulting in a total of 1,200 predicted peaks. The isolation list can be found in the supplementary materials.

parallel reaction monitoring mass spectrometry (PRM-MS) experiments were conducted at the John L. Holmes Mass Spectrometry Facility at the University of Ottawa with a Q Exactive™ Plus Hybrid Quadrupole-Orbitrap™ mass spectrometer, using the aforementioned isolation list to guide the scanning. The results were obtained from a single injection of a Thermo-Fisher Pierce™ HeLa protein digest standard. We opted to monitor for methylation in this sample because it is guaranteed to have a low missed tryptic cleavage rate (<10%) and minimal methionine oxidation and lysine carbamylation (<10%). Furthermore, these standards are thoroughly tested for quality, which improves the reproducibility of the results.

## Results and discussion

### MethylSight 2.0: performance and benchmarking

The predictive performances on the blind test set of the final models are tabulated in Table 4.

**Table 4.**
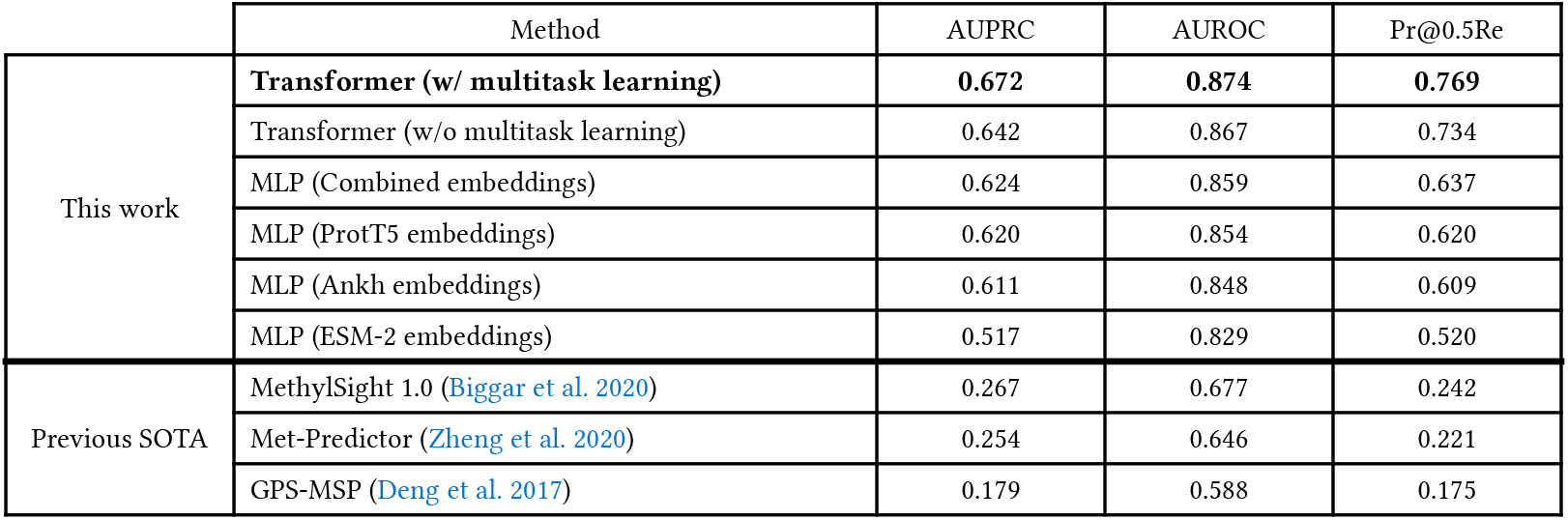
Prediction accuracy of our models and publicly available methods on the blind test set (1:6.5 imbalance)

|  | Method | AUPRC | AUROC | Pr@0.5Re |
| --- | --- | --- | --- | --- |
| This work | **Transformer (w/ multitask learning)** | **0.672** | **0.874** | **0.769** |
|  | Transformer (w/o multitask learning) | 0.642 | 0.867 | 0.734 |
|  | MLP (Combined embeddings) | 0.624 | 0.859 | 0.637 |
|  | MLP (ProtT5 embeddings) | 0.620 | 0.854 | 0.620 |
|  | MLP (Ankh embeddings) | 0.611 | 0.848 | 0.609 |
|  | MLP (ESM-2 embeddings) | 0.517 | 0.829 | 0.520 |
| Previous SOTA | MethylSight 1.0 (Biggar et al. 2020) | 0.267 | 0.677 | 0.242 |
|  | Met-Predictor (Zheng et al. 2020) | 0.254 | 0.646 | 0.221 |
|  | GPS-MSP (Deng et al. 2017) | 0.179 | 0.588 | 0.175 |

All models trained as part of this work perform significantly better on the blind test set than the previous state of the art. Indeed, our models outperformed GPS-MSP (Deng et al. 2017), MethylSight 1.0 (Biggar et al. 2020), and Met-Predictor (Zheng et al. 2020) in spite of the fact that these methods have likely encountered some of the sites in our test set during training while our model did not. In fact, our worst performing model, a MLP using lysine embeddings produce by the ESM-2 pLM was associated with a 25% improvement over the state of the art (SOTA) in terms of both AUPRC and precision at 0.5 recall (Pr@0.5Re).

Among the MLP models we trained on the four representations produced by the three pLMs considered, the model trained on ESM-2 embeddings performed noticeably worse relative to the other three representations which produced similar results, though the model trained on the combined embeddings appears to have a performance modestly superior to that of the MLPs trained on ProtT5 and Ankh embeddings. The relative performance of the different representations is consistent with the sizes of the pLMs, ProtT5 being 3 times the size of Ankh, and 4.6 times the size of ESM-2, in terms of learnable parameter counts. This observation is consistent with evidence that pLMs performance scales with model size following a power law (Cheng et al. 2024; Fournier et al. 2024).

The use of a transformer architecture trained “from scratch”specifically for the task of predicting lysine methylation prediction did allow for an improvement in performance over the use of lysine embeddings generated by all three foundational pLMs in MLPs. Our best single-task transformer model slightly outperformed the best MLP (*i*.*e*. the MLP trained on concatenated ProtT5-Ankh-ESM-2 embeddings), using AUPRC as a metric. However, it produced a more significant improvement in precision at the a 50% recall threshold of nearly 10%. This showcases the power of the self-attention mechanism, as attention layer parameters in our transformer model were learned specifically for the lysine methylation prediction task as opposed to the more general masked language modeling objective, as is was case for the foundational models.

Looking at the PRCs (PRCs) assessing model performance on our blind test set (Figure 4A), we see that implementing a multitask learning strategy leveraging knowledge about other PTMs is useful, as our best model (hyperparameters tabulated in Table 5) outperforms all other models over the entire range of possible recall values (or operating thresholds). However, this advantage is anticipated to dissipate at higher recall values assuming that the true class imbalance of methylation sites to non-methylation sites in the proteome is higher than that in the test set (*e*.*g*., 1:100 as opposed to 1:6.5; Figure 4C). This suggests that knowledge of other PTMs of lysines can indeed transfer to lysine methylation. This is consistent with our initial hypothesis as well as with the establish phenomenon of PTM competition wherein lysine modifying enzyme “compete” to modify specific lysine residues (Lee et al. 2023; Leutert, Entwisle, and Villén 2021; Shukri et al. 2023).

**Table 5.** Hyperparameters used to train the most accurate model (MethylSight 2.0)

| Parameter | Value |
| --- | --- |
| Learning rate | $8 \times 10^{-7}$ |
| Number of epochs | 100 |
| Weight factor of methylation loss ($\gamma$) | 20 |
| Batch size | 128 |
| Number of transformer blocks | 2 |
| Heads per self-attention layer | 4 |
| Width of the first (pre-attention) dense layer | 1,600 |
| Widths of the dense layers (classification heads) | 1,797; 1,803; 338; 493 |
| Embedding dimension | 1,024 |
| Dropout rate | 0.15 |

**Figure 4.**
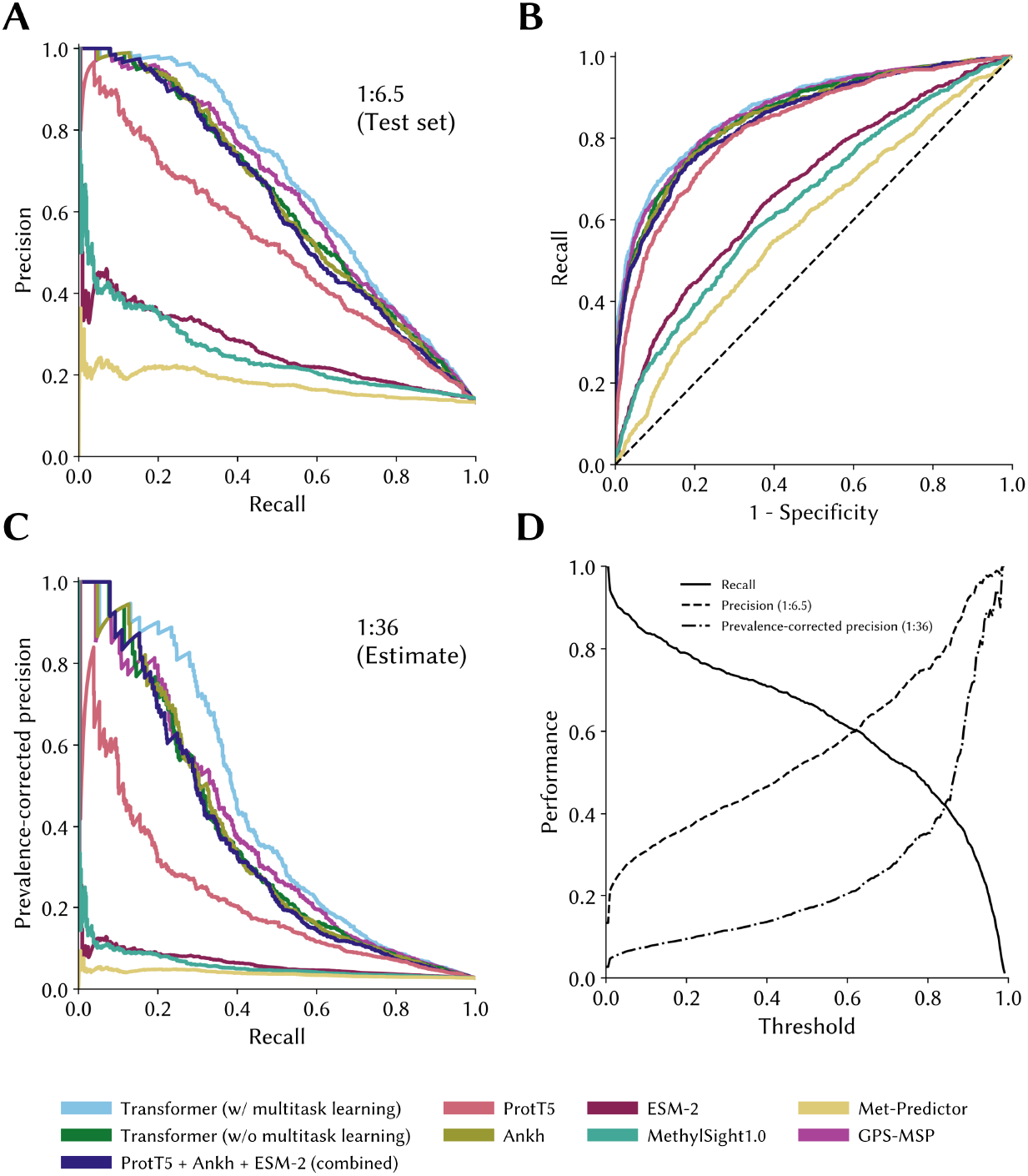
Precision-recall curves and receiver operating characteristic curves of the models on the independent test set. The precision-recall and receiver operating characteristic curves for the best performing models and/or training strategies were computed using a blind and independent test set of potential lysine methylation sites not seen during training. **(A)** The precision-recall curves are shown for the test set (1:6.5 imbalance). **(B)** The associated ROC curves are shown. **(C)** Prevalence-corrected precision assuming a true 1:36 imbalance between methylated and unmethylated sites provides more pessimistic estimate of performance. **(D)** The performance metrics are show for two different imbalance ratios (1:6.5 and 1:36).

### Identification and validation of novel lysine methylation sites

The PRM-MS experiments on a HeLa cell lysate guided with an isolation list listing tryptic peptides corresponding to MethylSight 2.0-predicted methylation sites revealed a significant number of hits (Figure 5A). In fact, 68 of the 100 sites predicted to be methylated produced transitions consistent with methylated peptides with a fair or better level of confidence. In contrast, for only 6 of the sites could evidence of the unmethylated peptide and no evidence of methylation be found. The results were inconclusive for 26 peptides, *i*.*e*. the peptide could not be detected, neither in a unmethylated state, nor in a methylated state. In the worst case where we consider all inconclusive sites to be negative, MethylSight 2.0 would achieve a precision of 68%. The precision becomes 91.9% if we discard sites for which no transitions could be detected from the analysis. It is likely that the precision we would have observed, if all results had been conclusive, would lie somewhere within that range. This suggests that an imbalance ratio of 1:36 is a reasonable estimate.

**Figure 5.**
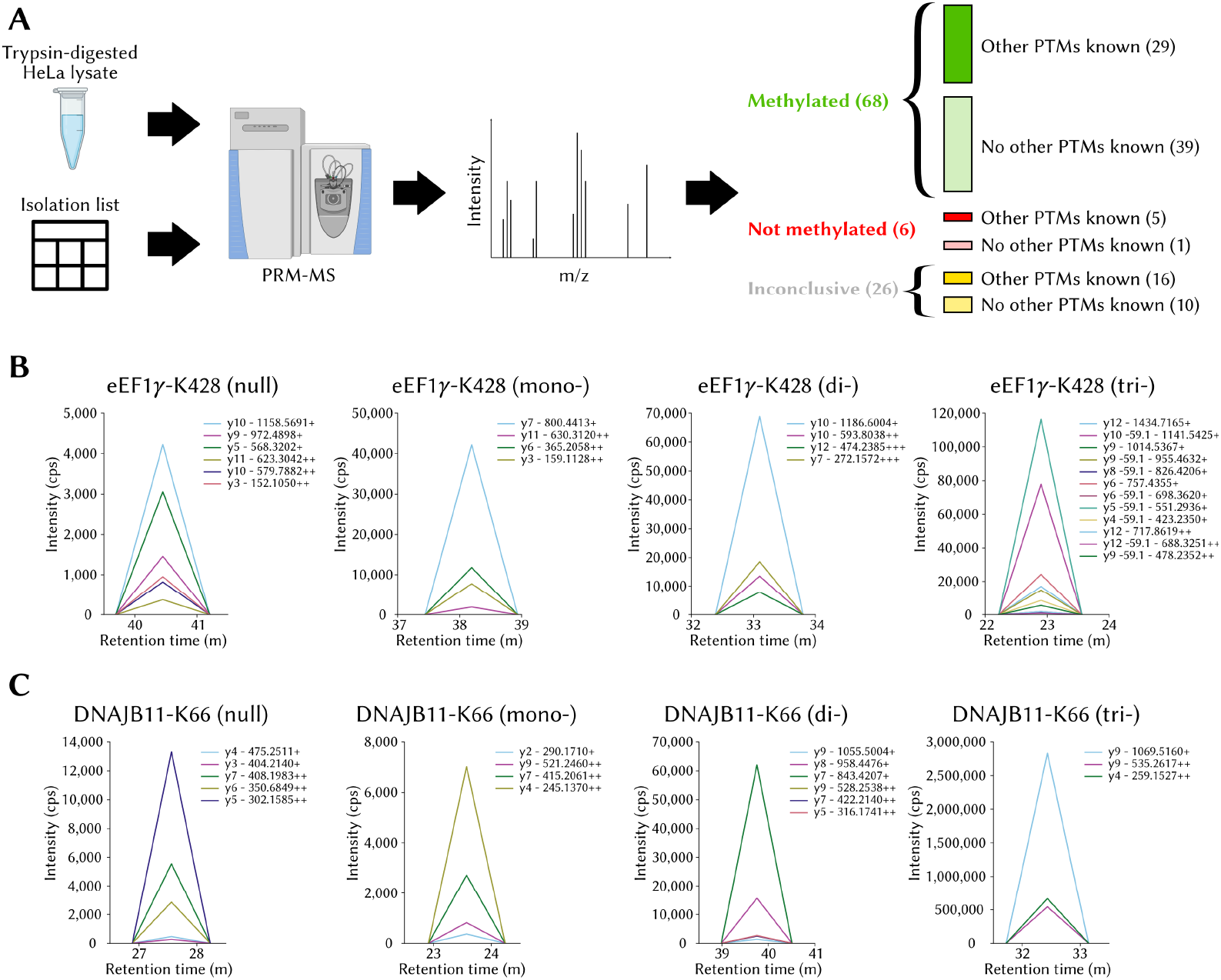
MethylSight 2.0-enabled discovery of novel lysine methylation sites with PRM-MS. **(A)** High-level overview of the methodology employed to validate 100 methylation sites identified with MethylSight 2.0 and distribution of the compiled results. **(B)** Transitions for the tryptic peptide containing eEF1*γ*-K428 (EYFSWEGAFQHVGK). The mass-to-charge ratios of the y ions are shown. The transitions of the null, mono-, di-, and tri-methylation states are plotted separately for clarity, because the measured intensity varies in scale. **(C)** Transitions for the tryptic peptide containing DNAJB11-K66 (`NPDDPQAQEK`).

Interestingly, we found that more methylation sites were found for lysines that were not known to be otherwise modified (39) than were found for lysines known to be acetylated, ubiquitinated, or sumoylated (29). This observation contradicts our initial hypothesis that more methylation sites would be detected among proteins which are known to be otherwise modified, because of their “modifiable” character. One plausible explanation is that one or more of these other modifications might have in fact competed with methylation, thus reducing its abundance and preventing its detection.

We choose here to highlight two sites occuring within proteins of high biological and clinical significance which produced transitions unequivocally consistent with methylation: the *γ* subunit of the Eukaryotic elongation factor 1 (eEF1*γ*) and DnaJ homolog subfamily B member 11 (DNAJB11).

eEF1*γ* is one of four subunit of the eEF1 complex, along with the *α, β* and *δ* subunits (Sasikumar, Perez, and Kinzy 2012). Though not believed to be a catalytically active member of the complex (Olarewaju et al. 2004), eEF1*γ* is believed to act as a structural scafford for the *α* subunit and to facilitate the complex’s function of bringing aminoacyl tRNAs to the ribosome for translation (Sasikumar, Perez, and Kinzy 2012). Beyond its role in the elongation factor 1 complex, eEF1*γ* is predicted to have “moonlighting” roles and be involved in several other biological processes including viral ribonucleic acid (RNA) transcription, oxidative stress response, cytoskeleton-membrane linking, and cellular trafficking (Negrutskii et al. 2023). MethylSight 2.0 correctly predicted the methylation of K428 in eEF1*γ* (Figure 5B), which is located within the C-terminus domain of the protein. While the N-terminus end of eEF1*γ* is known to interact with eEF1*α* and anchor it into the complex, the role of the C-terminus end of the protein is not a clearly understood, aside from the fact that it is highly conserved and protease resistant (Vanwetswinkel et al. 2003). There is some evidence that interaction with the *β* subunit of the eEF1 complex may occur at the C-terminus domain (Achilonu et al. 2018). Therefore, it is plausible that the methylation status of eEF1*γ*-K428 could modulate the interaction between these the *β* and *γ* subunits. Given that eEF1*γ* has been found to interact with actin (Olatona et al. 2024), it is possible that methylation of K428 could modulate this interaction and influence cytoskeleton dynamics if it indeed occurs at the C-terminus. eEF1*γ*’s clinical significance is supported by the observation that it overexpressed in gastric carcinoma (Mimori et al. 1995), colon adenocarcinoma (Chi, Jones, and Frazier 1992), and pancreatic cancer (Lew et al. 1992), in all likelihood so that cancer cells can satisfy the higher translation load required to adapt and proliferate. Altogether, our observation that eEF1*γ* is methylated on K438 combined with its involvement in key biological processes and cancer warrants further investigations into the biological significance of the modification.

Clear transitions consistent with the presence of a methyllysine were also recorded for the tryptic peptide fragment from DNAJB11 containing K66 (Figure 5C). DNAJB11 is a member of the DNAJ (or HSP40) subclass of family of heat shock proteins which all share a J-domain. The role of this highly conserved domain is to stimulate the hydrolysis of ATP by chaperones in the HSP70 protein family whose main function is stabilize or restore the native protein conformation of potentially misfolded client proteins under cellular stress (Kim and Hong 2022). Proteins in the DNAJ family have been implicated in tumor progression and metastasis (Kim and Hong 2022). DNAJB11 in particular has been overexpressed in pancreatic cancer, where exosomal DNAJB11 regulates expression of EGFR activates the MAPK pathway (Liu et al. 2022) and in liver cancer, by preventing alpha-1-antitrypsin degradation (Pan, Cao, and Gong 2018). In contrast, low DNAJB11 messenger RNA (mRNA) levels appear to be correlated with worse outcomes in thyroid carcinoma (Sun et al. 2021). In addition to it role in several cancers, research has shown that phosphorylation of T188 in DNAJB11 can reduce the aggregation of *α*-synuclein in Parkinson’s disease (Chen et al. 2024). As such, an association between K66 methylation and Parkinson’s disease is possible, either directly via an unknown mechanism, or indirectly, through the modulation of T188 phosphorylation via cross-talk, for example. In all cases, since it is located within the J-domain, it is likely that the methylation of K66 is biologically significant, and this result provides a rationale for the characterization of K66.

### The predicted human lysine methylome

Using conservative settings (*i*.*e*. at PCPr_1:36_ = 0.75), MethylSight 2.0 identified a total of 62,567 lysine methylation sites within 13,791 different proteins in the human proteome (Figure 6A). Based on our performance assessment and an estimated imbalance ratio of 1:36, we anticipate that out of these predicted sites, ∼47,000 (75%) are actual methylation sites. This figure, alone, is significantly higher than the ∼30,000 sites predicted with 63% precision by MethylSight 1.0. Given that the estimated recall of MethylSight 2.0 under these conditions is ∼30%, we estimate the size of the lysine methylome at ∼155,000.

**Figure 6.**
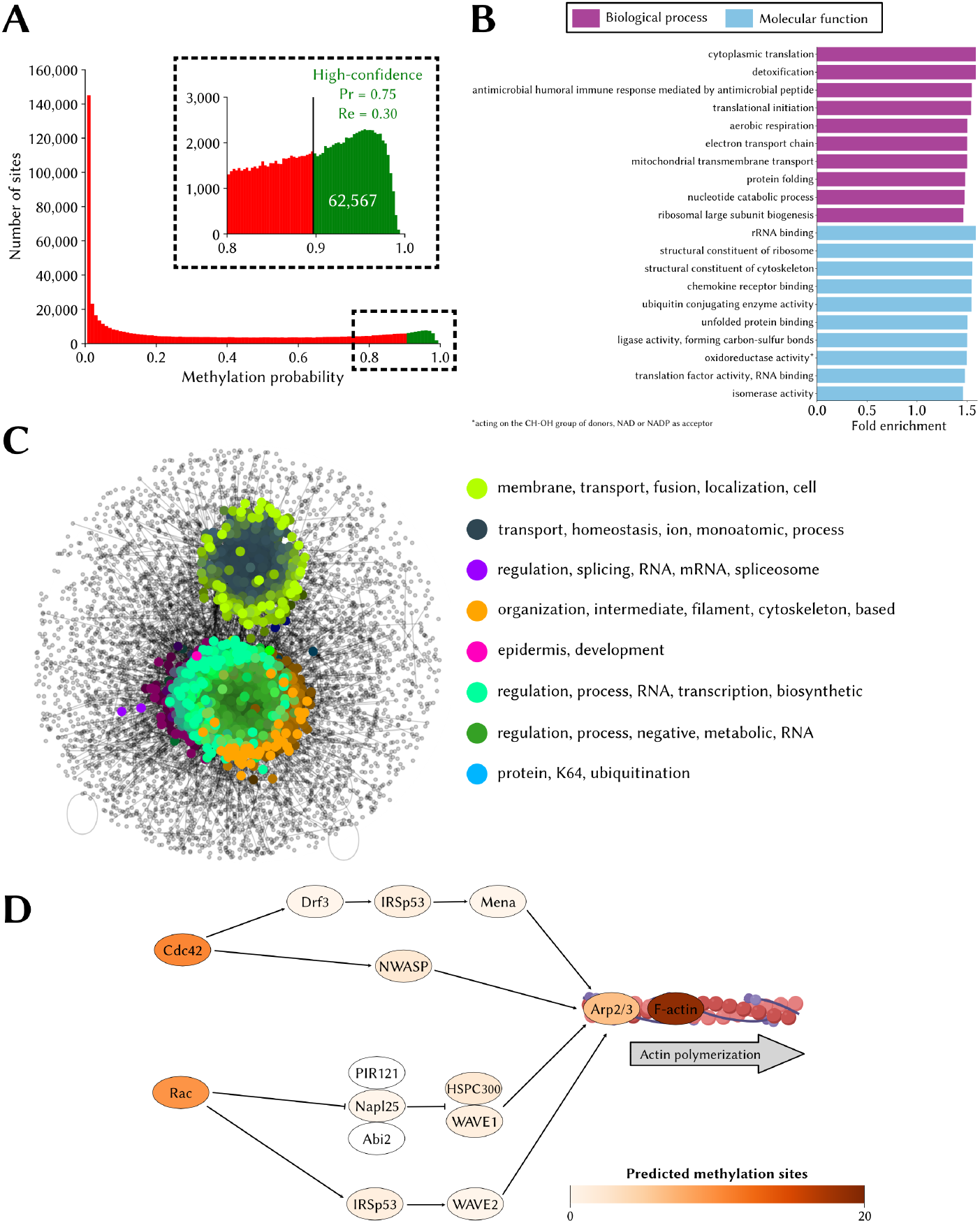
The lysine methylome as predicted by MethylSight 2.0. **(A)** Distribution of methylation probability for all 654,185 lysines in the human proteome. The sites predicted to be methylated *only* while operating under conservative settings (PCPr = 0.75) are shown in green. **(B)** Top 10 overrepresented gene ontology (GO) terms for the biological process (purple) and molecular function (blue) categories. Overrepresentation is statistically significant (*p*-value < 0.05; Fisher’s exact test with Bonferroni correction for multiple testing). **(C)** Visual representation of the spatial analysis of functional enrichment (SAFE) analysis results; *i*.*e*. functional domains within the interaction network of methylated proteins and the most frequent words present in the GO terms associated with proteins in the domains. **(D)** Subset of the actin cytoskeleton regulation pathway (KEGG: hsa04810). Proteins are colored on a white-to-red scale, with darker shades indicating a higher degree of methylation.

Statistically significant enrichment in several biological process and molecular function GO terms were identified (Figure 6B). Of note, we observed overrepresentation in methylated proteins of terms related to translation, ribosomal biogenesis and structure, and cytoskeleton structure. Enrichment of related terms in methylated proteins was also observed within the MethylSight 1.0-predicted lysine methylome (Biggar et al. 2020), and is also supported by the literature. For instance, methylation of several lysine residues in the human elongation factor 1A (eEF1A) is known to regulate ribosome biogenesis and actin cytoskeleton dynamics, among others (Hamey et al. 2017). The methylation of elongation factor eEF2 by the lysine methyltransferase (KMT) FAM86A is also known to regulate translation dynamics (Francis et al. 2024). The literature also supports the involvement of lysine methylation in cytoskeleton regulation. The role of lysine methylation in cytoskeleton regulation is well-established (Michail et al. 2025). The α-tubulin cytoskeletal protein is known to be tri-methylated by SETD2 (and acetylated) on K40, and loss of methylation has been associated with “catastrophic microtubule defects” which impair DNA repair mechanisms (Park et al. 2016) and cell cycle progression (Li and Li 2021). Recently, methylation of BCAR3 on K334 by SMYD2 in breast cancer was shown to enhance lamellipodia dynamics of breast cancer cells through the recruitment of Formin-like proteins which accelerate actin polymerization and facilitate cell proliferation and metastasis *in vivo* (Casanova et al. 2024).

Interestingly, our SAFE analysis of methylated proteins mapped onto the HuRI human interactome (Figure 6C) also shows subnetworks where overrepresentation of GO terms related to RNA processing and regulation and cytoskeleton organization is observed.

In Figure 6D, we illustrate the predicted prevalence of lysine methylation events in a subset of the actin cytoskeleton pathway (KEGG: hsa04810). Most proteins in this important subset of the pathway contain several lysine methylation sites.

### Cancer mutations associated with a predicted loss of methylation at a proximal site

MethylSight 2.0 was used to predict the impact of the 1,000 most frequent missense mutations in the COSMIC database (Sondka et al. 2024) on the predicted methylation score of lysines within the mutant protein. The COSMIC database contains a curated list of somatic mutations and their impact in cancer.

We found that scores were relatively insensitive to these mutations except in select cases where the mutations were in close proximity with a lysine. Interestingly, we only observed predicted *loss* of methylation (using the conservative operating threshold). We detected 25 lysines that were no longer predicted to be methylated and associated with a decrease in methylation score ≥ 0.02 in presence of a mutation (Figure 7A).

**Figure 7.**
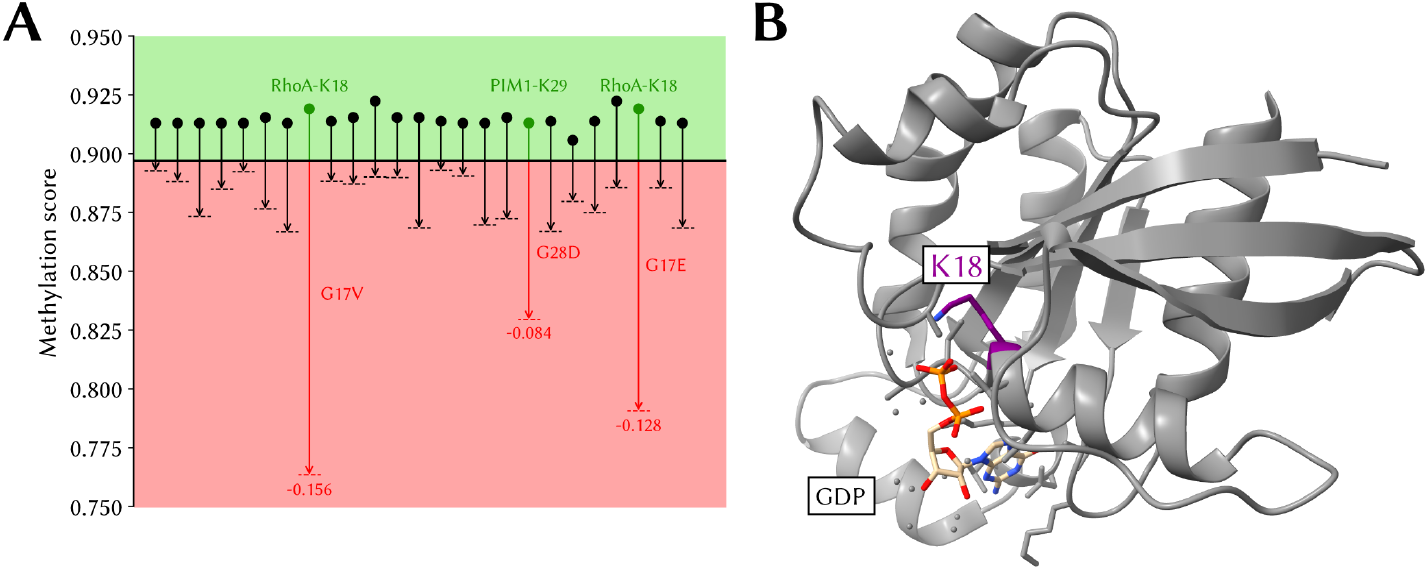
Predicted loss of methylation in oncogenic proteins. **(A)** Changes in predicted methylation score resulting in predicted loss of methylation associated with the 1,000 most commonly reported cancer-associated single amino acid mutations in the Catalog of Somatic Mutations reported in the Cancer (COSMIC) database. Shown are score changes of at least 2%. **(B)** Position of K18 (in purple) in the X-ray structure of RhoA (PDB: 1DPF). The GDP co-factor is in beige with the two phosphate groups in orange.

The most striking loss of predicted methylation are associated with the RhoA^G17V^ and RhoA^G17E^ mutations. In these mutants, the neighbouring K18 is no longer predicted to be a methylation site. K18, like the mutated G17, is located amidst the GDP binding pocket (Figure 7B).

While methylation of RhoA-K18 has never been – to the best of our knowledge – confirmed experimentally, it is possible that methylated K18 could modulate RhoA activity. In fact, though it is believed that mutations in G17 impair GDP/GTP binding (Sakata-Yanagimoto et al. 2014), it is not known exactly *how* this mutation impairs binding of GDP/GTP. Given the proximity of K18 to the GTP/GDP binding site, it is not implausible that alteration of the methylation status could alter RhoA’s ability to bind and release GTP/GDP or coordinate Mg^2+^.

Taken together, this provides an interesting avenue for further investigation so as to determine whether ❶ RhoA-K18 is a true methylation site and ❷ loss of methylation occurs in these mutants *in vitro*, and ❸ this loss of methylation directly impacts GTP/GDP binding.

### Performance of MethylSight 2.0 on non-human proteins

We deployed MethylSight 2.0 on the set of all known non-human lysine methylation sites catalogued in the PhosphoSitePlus database (360 sites). Interestingly, MethylSight 2.0 achieved a recall of 0.383 when applied to these sites (at PCPr_1:36_ = 0.75). This is on the same order as its predicted recall (*i*.*e*. 0.299) on human lysines operating at the same decision threshold. This suggests that methylation sites in non-human organisms share homology with sites in human proteins.

The observation that MethylSight 2.0 achieved a better recall than predicted on this set of sites provides some evidence that the imbalance between methylated and non-methylated lysines could actually be lower than the one we estimated (*i*.*e*. 1:36) at the proteome scale, but further experiments would be required to confirm this.

### The MethylSight 2.0 server

To make MethylSight 2.0 accessible to the broader community, we implemented a web server (Figure 8) which can be accessed at https://methylsight2.cu-bic.ca. The server is easy-to-use and allows users to select the operating threshold (precision and recall) that suits them best, depending on the application.

**Figure 8.**
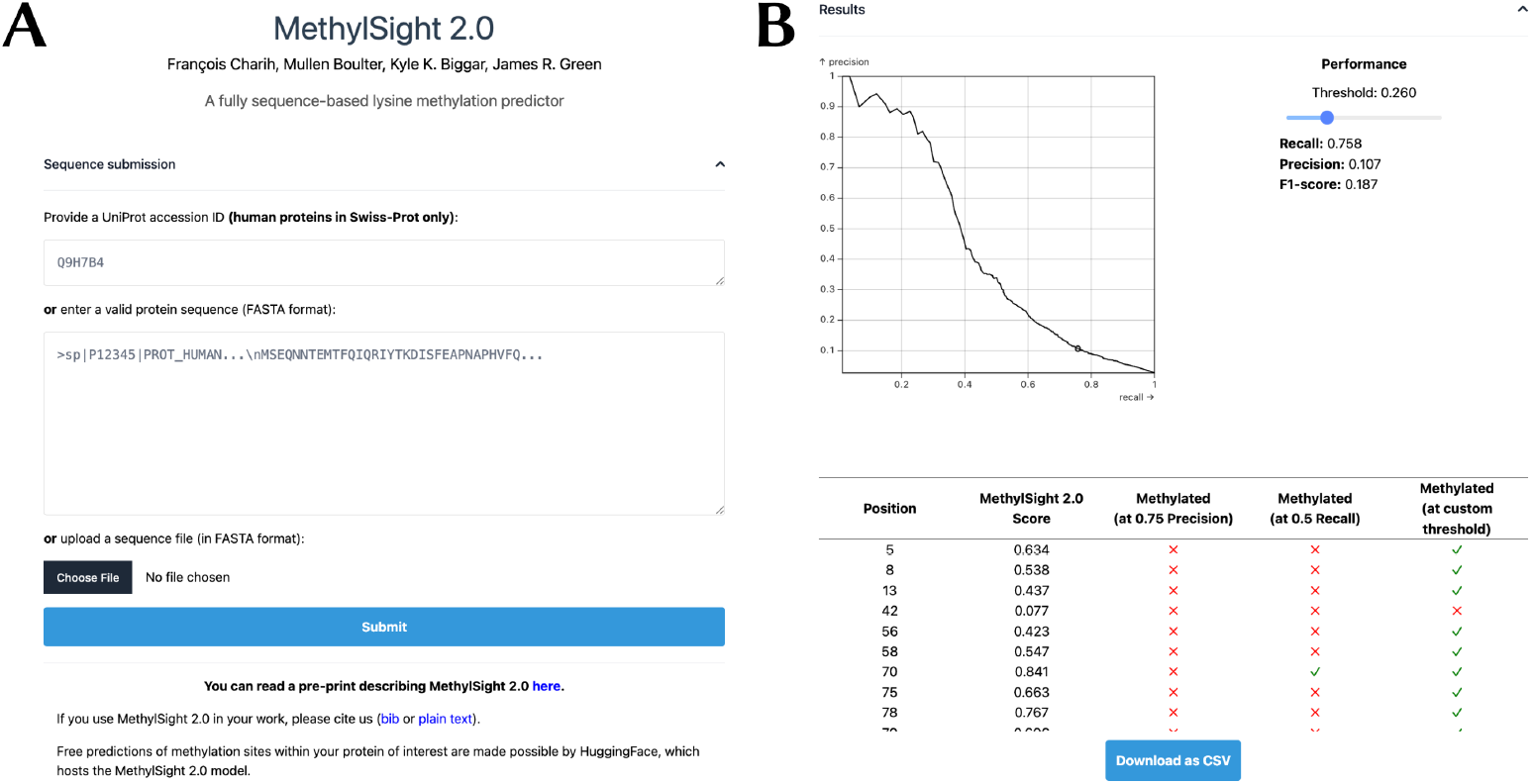
MethylSight 2.0 web server. **(A)** The user can either provide the UniProt accession ID or the sequence of the human protein of interest. If the protein is from another organism or is a non-canonical human protein (*e*.*g*., isoform or mutant), the user must provide a FASTA-formatted sequence through a text box or uploading the equivalent FASTA file. **(B)** Once the results are available, a precision-recall curve computed for MethylSight 2.0 on our test set with an assumed real imbalance ratio of 1:36 is presented. This provides the user with an visual interpretation of the operating threshold and the optimal threshold which can be tuned with a slider. The results are presented as a table which can be downloaded in CSV format.

The web server processes *individual* protein sequences. Users interested in batch predictions may run MethylSight 2.0 as a standalone software on their own hardware. The Methylsight 2.0 source code, model weights, and instructions on how to use the software can be found on GitHub: https://github.com/GreenCUBIC/MethylSight2.

## Conclusion

Our work demonstrates that deep representations learned by pLMs trained on tens of millions of protein are rich in information directly relevant for the task of lysine methylation site prediction and significantly improve the quality of the predictions. In fact, using these deep representations, we successfully trained model that achieved more than double the AUPRC of previous models trained on human-engineered descriptors of protein sequences, such as those generated by ProtDCal (Ruiz-Blanco et al. 2015) alongside the SVM-based MethylSight 1.0 predictor. We also showed that leveraging knowledge about other PTMs by means of a multitask learning strategy can further enhance the quality of the predictions. To the best of our knowledge, our model MethylSight 2.0 is the first lysine methylation prediction model to leverage pLM-generated representations and to employ a multitask learning strategy to extract knowledge from useful data that would otherwise be left unexploited.

In an effort to validate the predictions made by MethylSight 2.0 and show that it can successfully guide lysine methylation site discovery *in vivo*, we performed a validation PRM-MS experiment guided by MethylSight 2.0 predictions on a high-quality HeLa cell lysate. We uncovered 68 previously unidentified lysine methylation sites, some of which among proteins of high biological and/or therapeutic relevance, further showing the usefulness of our model as a drug target identification tool.

Applying MethylSight 2.0 to the human proteome provides insight into the extent of the lysine methylome. In fact, our analyses of the MethylSight 2.0-predicted lysine methylome suggests that the number of methyllysines in the human proteome may be even larger than previously believed (∼50,000 as per Biggar et al. 2020), though this number remains difficult to estimate, given that our validation experiment was limited to 100 sites, and it seems unlikely that so few sites would be representative of the entire human proteome. Additional validation experiments would be required to get a more accurate portrait of the human lysine methylome landscape.

Lysine methylation is a highly dynamic process which competes with several other PTMs, and a given lysine may be methylated to varying degrees under different conditions, *e*.*g*., during development, in response to an environmental trigger, or in specific tissue types. Given that our model was trained in a tissue-blind fashion, *i*.*e*. positive sites in the training set were known to be methylated in *at least* one tissue type, we anticipate that validation experiments on methylation sites identified with MethylSight 2.0 may need to be performed in more than one cell type. There could be significant value in training a cell-specific lysine methylation predictor that could predict whether a lysine is methylated *in a given tissue type*, but training such a model would require tissue-specific datasets which are currently not publicly available.

Furthermore, in this study, we did not distinguish between the three possible methylation states. It is important to acknowledge that different methylation states can be associated with different – sometimes opposite – phenotypes. Several other groups have attempted to address this problem (Deng et al. 2017; Ju, Cao, and Gu 2015; Zheng et al. 2020), but with limited success. Further research is needed to design an accurate predictor of mono-, di-, and trimethylated lysines.

Finally, MethylSight 2.0 does not attempt to associate a KMT with sites predicted to be methylated. This challenge is of prime importance, as therapeutic intervention would normally target the KMT or lysine demethylase (KDM) responsible for the modification. However, it is non-trivial given the scarcity of data for some KMTs, which in certain cases only have a few dozen known substrates or fewer. At the time of writing, we are aware of only one KMT-specific model for SET8 (Biggar et al. 2025) which could be used in conjunction with MethylSight 2.0.

Taken together, this work constitutes a significant contribution toward the elucidation of the human lysine methylome. In addition, MethylSight 2.0 can be deployed in a targeted fashion to determine whether lysines within proteins involved in a pathway of interest are probable methylation site. It therefore affords experimentalists with a tool which can help formulate rational hypotheses, guide experiments, and cut costs through prioritization of candidates for validation experiments.

## Author contributions

**François Charih:** Conceptualization, Methodology, Software, Investigation, Formal analysis, Data Curation, Visualization, Writing - Original Draft, **Mullen Boulter:** Formal analysis, **Kyle K. Biggar:** Conceptualization, Resources, Formal analysis, Writing - Review & Editing, Funding acquisition, **James R. Green:** Conceptualization, Writing - Review & Editing, Funding acquisition

All authors approved of the manuscript.

## References

Achilonu, Ikechukwu, Nnenna Elebo, Babongiwe Hlabano, Gavin R. Owen, Maria Papathanasopoulos, and Heini W. Dirr. 2018. An Update on the Biophysical Character of the Human Eukaryotic Elongation Factor 1 Beta: Perspectives from Interaction with Elongation Factor 1 Gamma. Journal of Molecular Recognition 31 (7):e2708. doi:10.1002/jmr.2708.

Akiba, Takuya, Shotaro Sano, Toshihiko Yanase, Takeru Ohta, and Masanori Koyama. 2019. Optuna: A Next-generation Hyperparameter Optimization Framework. In Proceedings of the 25th ACM SIGKDD International Conference on Knowledge Discovery & Data Mining, 2623–2631. KDD ‘19. New York, NY, USA: Association for Computing Machinery. doi:10.1145/3292500.3330701.

Avraham, Orly, Tomer Tsaban, Ziv Ben-Aharon, Linoy Tsaban, and Ora Schueler-Furman. 2023. Protein Language Models Can Capture Protein Quaternary State. BMC Bioinformatics 24 (1):433. doi:10.1186/s12859-023-05549-w.

Bepler, Tristan, and Bonnie Berger. 2021. Learning the Protein Language: Evolution, Structure, And Function. Cell Systems 12 (6):654–669. doi:10.1016/j.cels.2021.05.017.

Berryhill, Christine A., Jocelyne N. Hanquier, Emma H. Doud, Eric Cordeiro-Spinetti, Bradley M. Dickson, Scott B. Rothbart, Amber L. Mosley, and Evan M. Cornett. 2023. Global Lysine Methylome Profiling Using Systematically Characterized Affinity Reagents. Scientific Reports 13 (1):377. doi:10.1038/s41598-022-27175-x.

Bhat, Suhaas, Kalyan Palepu, Lauren Hong, Joey Mao, Tianzheng Ye, Rema Iyer, Lin Zhao, Tianlai Chen, Sophia Vincoff, Rio Watson, et al. 2025. De Novo Design of Peptide Binders to Conformationally Diverse Targets with Contrastive Language Modeling. Science Advances 11 (4). American Association for the Advancement of Science:eadr8638. doi:10.1126/sciadv.adr8638.

Biggar, Kyle K., and Shawn S.-C. Li. 2015. Non-Histone Protein Methylation as a Regulator of Cellular Signalling and Function. Nature Reviews Molecular Cell Biology 16 (1):5–17. doi:10.1038/nrm3915.

Biggar, Kyle K., Francois Charih, Huadong Liu, Yasser B. Ruiz-Blanco, Leanne Stalker, Anand Chopra, Justin Connolly, Hemanta Adhikary, Kristin Frensemier, Matthew Hoekstra, et al. 2020. Proteome-Wide Prediction of Lysine Methylation Leads to Identification of H2BK43 Methylation and Outlines the Potential Methyllysine Proteome. Cell Reports 32 (2):107896. doi:10.1016/j.celrep.2020.107896.

Biggar, Kyle, Nashira Ridgeway, Anand Chopra, Valentina Lukinovic, Michal Feldman, Francois Charih, Dan Levy, and James Green. 2025. Machine Learning-Based Exploration of Enzyme-Substrate Networks: SET8-mediated Methyllysine and Its Changing Impact within Cancer Proteomes. Communications Chemistry (Accepted).

Brixi, Garyk, Tianzheng Ye, Lauren Hong, Tian Wang, Connor Monticello, Natalia Lopez-Barbosa, Sophia Vincoff, Vivian Yudistyra, Lin Zhao, Elena Haarer, et al. 2023. SaLT&PepPr Is an Interface-Predicting Language Model for Designing Peptide-Guided Protein Degraders. Communications Biology 6 (1):1081. doi:10.1038/s42003-023-05464-z.

Carlson, Scott M., and Or Gozani. 2016. Nonhistone Lysine Methylation in the Regulation of Cancer Pathways. Cold Spring Harbor Perspectives in Medicine 6 (11):a26435. doi:10.1101/cshperspect.a026435.

Casanova, Alexandre G., Gael S. Roth, Simone Hausmann, Xiaoyin Lu, Ludivine J. M. Bischoff, Emilie M. Froeliger, Lucid Belmudes, Ekaterina Bourova-Flin, Natasha M. Flores, Ana Morales Benitez, et al. 2024. Cytoskeleton Remodeling Induced by SMYD2 Methyltransferase Drives Breast Cancer Metastasis. Cell Discovery 10 (1). Nature Publishing Group:1–22. doi:10.1038/s41421-023-00644-x.

Chen, Hu, Yu Xue, Ni Huang, Xuebiao Yao, and Zhirong Sun. 2006. MeMo: A Web Tool for Prediction of Protein Methylation Modifications. Nucleic Acids Research 34 (suppl_2):W249–W253. doi:10.1093/nar/gkl233.

Chen, Huan-Yun, Chia-Yu Liao, Hsun Li, Yi-Ci Ke, Chin-Hsien Lin, and Shu-Chun Teng. 2024. ATM-mediated Co-Chaperone DNAJB11 Phosphorylation Facilitates A-Synuclein Folding Upon DNA Double-Stranded Breaks. NAR Molecular Medicine 1 (2):ugae7. doi:10.1093/narmme/ugae007.

Chen, Leo Tianlai, Zachary Quinn, Madeleine Dumas, Christina Peng, Lauren Hong, Moises Lopez-Gonzalez, Alexander Mestre, Rio Watson, Sophia Vincoff, Lin Zhao, et al. 2025. Target Sequence-Conditioned Design of Peptide Binders Using Masked Language Modeling. Nature Biotechnology, August. Nature Publishing Group, 1–9. doi:10.1038/s41587-025-02761-2.

Cheng, Xingyi, Bo Chen, Pan Li, Jing Gong, Jie Tang, and Le Song. 2024. Training Compute-Optimal Protein Language Models. Preprint: bioRxiv. doi:10.1101/2024.06.06.597716.

Chi, K., D. V. Jones, and M. L. Frazier. 1992. Expression of an Elongation Factor 1 Gamma-Related Sequence in Adenocarcinomas of the Colon. Gastroenterology 103 (1):98–102. doi:10.1016/0016-5085(92)91101-9.

Damgaard, Rune Busk. 2021. The Ubiquitin System: From Cell Signalling to Disease Biology and New Therapeutic Opportunities. Cell Death & Differentiation 28 (2). Nature Publishing Group:423–426. doi:10.1038/s41418-020-00703-w.

Deng, Wankun, Yongbo Wang, Lili Ma, Ying Zhang, Shahid Ullah, and Yu Xue. 2017. Computational Prediction of Methylation Types of Covalently Modified Lysine and Arginine Residues in Proteins. Briefings in Bioinformatics 18 (4):647–658. doi:10.1093/bib/bbw041.

Elnaggar, Ahmed, Hazem Essam, Wafaa Salah-Eldin, Walid Moustafa, Mohamed Elkerdawy, Charlotte Rochereau, and Burkhard Rost. 2023. Ankh: Optimized Protein Language Model Unlocks General-Purpose Modelling. Preprint: bioRxiv. doi:10.1101/2023.01.16.524265.

Elnaggar, Ahmed, Michael Heinzinger, Christian Dallago, Ghalia Rehawi, Yu Wang, Llion Jones, Tom Gibbs, Tamas Feher, Christoph Angerer, Martin Steinegger, et al. 2021. ProtTrans: Towards Cracking the Language of Lifes Code Through Self-Supervised Deep Learning and High Performance Computing. IEEE Transactions on Pattern Analysis and Machine Intelligence, 1. doi:10.1109/TPAMI.2021.3095381.

Feoli, Alessandra, Monica Viviano, Alessandra Cipriano, Ciro Milite, Sabrina Castellano, and Gianluca Sbardella. 2022. Lysine Methyltransferase Inhibitors: Where We Are Now. RSC Chemical Biology 3 (4):359–406. doi:10.1039/D1CB00196E.

Fournier, Quentin, Robert M. Vernon, Almer Van Der Sloot, Benjamin Schulz, Sarath Chandar, and Christopher James Langmead. 2024. Protein Language Models: Is Scaling Necessary?. Preprint: bioRxiv. doi:10.1101/2024.09.23.614603.

Francis, Joel William, Simone Hausmann, Sabeen Ikram, Kunlun Yin, Robert Mealey-Farr, Natasha Mahealani Flores, Annie Truc Trinh, Tourkian Chasan, Julia Thompson, Pawel Karol Mazur, et al. 2024. FAM86A Methylation of eEF2 Links mRNA Translation Elongation to Tumorigenesis. Molecular Cell 84 (9):1753–1763. doi:10.1016/j.molcel.2024.02.037.

Fu, Limin, Beifang Niu, Zhengwei Zhu, Sitao Wu, and Weizhong Li. 2012. CD-HIT: Accelerated for Clustering the Next-Generation Sequencing Data. Bioinformatics 28 (23):3150–3152. doi:10.1093/bioinformatics/bts565.

Geiss-Friedlander, Ruth, and Frauke Melchior. 2007. Concepts in Sumoylation: A Decade on. Nature Reviews Molecular Cell Biology 8 (12):947–956. doi:10.1038/nrm2293.

Hamey, Joshua J., Beeke Wienert, Kate G. R. Quinlan, and Marc R. Wilkins. 2017. METTL21B Is a Novel Human Lysine Methyltransferase of Translation Elongation Factor 1A: Discovery by CRISPR/Cas9 Knockout. Molecular & Cellular Proteomics 16 (12):2229–2242. doi:10.1074/mcp.M116.066308.

Han, Dong, Mengxi Huang, Ting Wang, Zhiping Li, Yanyan Chen, Chao Liu, Zengjie Lei, and Xiaoyuan Chu. 2019. Lysine Methylation of Transcription Factors in Cancer. Cell Death & Disease 10 (4):290. doi:10.1038/s41419-019-1524-2.

Hao, Xiaohu, and Long Fan. 2024. ProtT5 and Random Forests-Based Viscosity Prediction Method for Therapeutic mAbs. European Journal of Pharmaceutical Sciences 194 (March):106705. doi:10.1016/j.ejps.2024.106705.

Hornbeck, Peter V., Bin Zhang, Beth Murray, Jon M. Kornhauser, Vaughan Latham, and Elzbieta Skrzypek. 2015. PhosphoSitePlus, 2014: Mutations, PTMs and Recalibrations. Nucleic Acids Research 43 (Database issue):D512–520. doi:10.1093/nar/gku1267.

Huang, Mingyao, Zirong Jiang, Yadan Xu, Chaoshen Wu, Wei Ding, Xuli Meng, and Da Qian. 2024. Methylation Modification of Non-Histone Proteins in Breast Cancer: An Emerging Targeted Therapeutic Strategy. Pharmacological Research 208 (October):107354. doi:10.1016/j.phrs.2024.107354.

Høie Magnus Haraldson, Erik Nicolas Kiehl, Bent Petersen, Morten Nielsen, Ole Winther, Henrik Nielsen, Jeppe Hallgren, and Paolo Marcatili. 2022. NetSurfP-3.0: Accurate and Fast Prediction of Protein Structural Features by Protein Language Models and Deep Learning. Nucleic Acids Research 50 (W1):W510–W515. doi:10.1093/nar/gkac439.

Ju, Zhe, Jun-Zhe Cao, and Hong Gu. 2015. iLM-2L: A Two-Level Predictor for Identifying Protein Lysine Methylation Sites and Their Methylation Degrees by Incorporating K-gap Amino Acid Pairs into Chou’s General PseAAC. Journal of Theoretical Biology 385 (November):50–57. doi:10.1016/j.jtbi.2015.07.030.

Kawashima, Shuichi, Piotr Pokarowski, Maria Pokarowska, Andrzej Kolinski, Toshiaki Katayama, and Minoru Kanehisa. 2008. AAindex: Amino Acid Index Database, Progress Report 2008. Nucleic Acids Research 36 (Database issue):D202–205. doi:10.1093/nar/gkm998.

Kim, Eunhee, Misuk Kim, Dong-Hun Woo, Yongjae Shin, Jihye Shin, Nakho Chang, Young Taek Oh, Hong Kim, Jingeun Rheey, Ichiro Nakano, et al. 2013. Phosphorylation of EZH2 Activates STAT3 Signaling via STAT3 Methylation and Promotes Tumorigenicity of Glioblastoma Stem-Like Cells. Cancer Cell 23 (6):839–852. doi:10.1016/j.ccr.2013.04.008.

Kim, Hye-Youn, and Suntaek Hong. 2022. Multi-Faceted Roles of DNAJB Protein in Cancer Metastasis and Clinical Implications. International Journal of Molecular Sciences 23 (23):14970. doi:10.3390/ijms232314970.

Kingma, Diederik P., and Jimmy Ba. 2017. Adam: A Method for Stochastic Optimization. Preprint: arXiv. doi:10.48550/arXiv.1412.6980.

Lanouette, Sylvain, Vanessa Mongeon, Daniel Figeys, and Jean-François Couture. 2014. The Functional Diversity of Protein Lysine Methylation. Molecular Systems Biology 10 (4):724. doi:10.1002/msb.134974.

Lee, Ji Min, Henrik M. Hammarén, Mikhail M. Savitski, and Sung Hee Baek. 2023. Control of Protein Stability by Post-Translational Modifications. Nature Communications 14 (1). Nature Publishing Group:201. doi:10.1038/s41467-023-35795-8.

Leutert, Mario, Samuel W. Entwisle, and Judit Villén. 2021. Decoding Post-Translational Modification Crosstalk With Proteomics. Molecular & Cellular Proteomics : MCP 20 (July):100129. doi:10.1016/j.mcpro.2021.100129.

Levitsky, Lev I., Joshua A. Klein, Mark V. Ivanov, and Mikhail V. Gorshkov. 2018. Pyteomics 4.0: Five Years of Development of a Python Proteomics Framework. Journal of Proteome Research 18 (2). American Chemical Society:709–714. doi:10.1021/acs.jproteome.8b00717.

Lew, Y., D. V. Jones, W. M. Mars, D. Evans, D. Byrd, and M. L. Frazier. 1992. Expression of Elongation Factor-1 Gamma-Related Sequence in Human Pancreatic Cancer. Pancreas 7 (2):144–152. doi:10.1097/00006676-199203000-00003.

Li, Ang, Yingwei Deng, Yan Tan, and Min Chen. 2021. A Transfer Learning-Based Approach for Lysine Propionylation Prediction. Frontiers in Physiology 12 (April). Frontiers. doi:10.3389/fphys.2021.658633.

Li, Linda Xiaoyan, and Xiaogang Li. 2021. Epigenetically Mediated Ciliogenesis and Cell Cycle Regulation, And Their Translational Potential. Cells 10 (7). Multidisciplinary Digital Publishing Institute:1662. doi:10.3390/cells10071662.

Lin, Zeming, Halil Akin, Roshan Rao, Brian Hie, Zhongkai Zhu, Wenting Lu, Nikita Smetanin, Robert Verkuil, Ori Kabeli, Yaniv Shmueli, et al. 2023. Evolutionary-Scale Prediction of Atomic-Level Protein Structure with a Language Model. Science 379 (6637). American Association for the Advancement of Science:1123–1130. doi:10.1126/science.ade2574.

Liu, Peng, Fuqiang Zu, Hui Chen, Xiaoli Yin, and Xiaodong Tan. 2022. Exosomal DNAJB11 Promotes the Development of Pancreatic Cancer by Modulating the EGFR/MAPK Pathway. Cellular & Molecular Biology Letters 27 (October):87. doi:10.1186/s11658-022-00390-0.

Lukinović, Valentina, Alexandre G. Casanova, Gael S. Roth, Florent Chuffart, and Nicolas Reynoird. 2020. Lysine Methyltransferases Signaling: Histones Are Just the Tip of the Iceberg. Current Protein and Peptide Science 21 (7):655–674. doi:10.2174/1871527319666200102101608.

Michail, Christina, Fernando Rodrigues Lima, Mireille Viguier, and Frédérique Deshayes. 2025. Structure and Function of the Lysine Methyltransferase SETD2 in Cancer: From Histones to Cytoskeleton. Neoplasia 59 (January):101090. doi:10.1016/j.neo.2024.101090.

Mimori, K., M. Mori, S. Tanaka, T. Akiyoshi, and K. Sugimachi. 1995. The Overexpression of Elongation Factor 1 Gamma mRNA in Gastric Carcinoma. Cancer 75 (6Suppl):1446–1449. doi:10.1002/1097-0142(19950315)75:6+<1446::aid-cncr2820751509>3.0.co;2-p.

Narita, Takeo, Brian T. Weinert, and Chunaram Choudhary. 2019. Functions and Mechanisms of Non-Histone Protein Acetylation. Nature Reviews Molecular Cell Biology 20 (3):156–174. doi:10.1038/s41580-018-0081-3.

Negrutskii, B. S., V. F. Shalak, O. V. Novosylna, L. V. Porubleva, D. M. Lozhko, and A. V. El’skaya. 2023. The eEF1 Family of Mammalian Translation Elongation Factors. BBA Advances 3 (January):100067. doi:10.1016/j.bbadva.2022.100067.

Olarewaju, Olubunmi, Pedro A. Ortiz, Wasimul Q. Chowdhury, Ishita Chatterjee, and Terri Goss Kinzy. 2004. The Translation Elongation Factor eEF1B Plays a Role in the Oxidative Stress Response Pathway. RNA Biology 1 (2):89–94. doi:10.4161/rna.1.2.1033.

Olatona, Olusola A., Sayantan R. Choudhury, Ray Kresman, and Carol A. Heckman. 2024. Candidate Proteins Interacting with Cytoskeleton in Cells from the Basal Airway Epithelium in Vitro. Frontiers in Molecular Biosciences 11 (July):1423503. doi:10.3389/fmolb.2024.1423503.

Pan, Junjiang, Ding Cao, and Jianping Gong. 2018. The Endoplasmic Reticulum Co-Chaperone ERdj3/DNAJB11 Promotes Hepatocellular Carcinoma Progression Through Suppressing AATZ Degradation. Future Oncology 14 (29):3001–3013. doi:10.2217/fon-2018-0401.

Park, In Young, Reid T. Powell, Durga Nand Tripathi, Ruhee Dere, Thai H. Ho, T. Lynne Blasius, Yun-Chen Chiang, Ian J. Davis, Catherine C. Fahey, Kathryn E. Hacker, et al. 2016. Dual Chromatin and Cytoskeletal Remodeling by SETD2. Cell 166 (4):950–962. doi:10.1016/j.cell.2016.07.005.

Paszke, Adam, Sam Gross, Francisco Massa, Adam Lerer, James Bradbury, Gregory Chanan, Trevor Killeen, Zeming Lin, Natalia Gimelshein, Luca Antiga, et al. 2019. PyTorch: An Imperative Style, High-Performance Deep Learning Library. Preprint: arXiv. doi:10.48550/arXiv.1912.01703.

Peng, Fred Zhangzhi, Chentong Wang, Tong Chen, Benjamin Schussheim, Sophia Vincoff, and Pranam Chatterjee. 2025. PTM-Mamba: A PTM-aware Protein Language Model with Bidirectional Gated Mamba Blocks. Nature Methods 22 (5). Nature Publishing Group:945–949. doi:10.1038/s41592-025-02656-9.

Petersen, Bent, Thomas Petersen, Pernille Andersen, Morten Nielsen, and Claus Lundegaard. 2009. A Generic Method for Assignment of Reliability Scores Applied to Solvent Accessibility Predictions. BMC Structural Biology 9 (1):51. doi:10.1186/1472-6807-9-51.

Qiu, Wang-Ren, Xuan Xiao, Wei-Zhong Lin, and Kuo-Chen Chou. 2014. iMethyl-PseAAC: Identification of Protein Methylation Sites via a Pseudo Amino Acid Composition Approach. Biomed Research International 2014 (May):947416. doi:10.1155/2014/947416.

Ruiz-Blanco, Yasser B., Waldo Paz, James Green, and Yovani Marrero-Ponce. 2015. ProtDCal: A Program to Compute General-Purpose-Numerical Descriptors for Sequences and 3D-structures of Proteins. BMC Bioinformatics 16 (1):162. doi:10.1186/s12859-015-0586-0.

Sakata-Yanagimoto, Mamiko, Terukazu Enami, Kenichi Yoshida, Yuichi Shiraishi, Ryohei Ishii, Yasuyuki Miyake, Hideharu Muto, Naoko Tsuyama, Aiko Sato-Otsubo, Yusuke Okuno, et al. 2014. Somatic RHOA Mutation in Angioimmunoblastic T Cell Lymphoma. Nature Genetics 46 (2):171–175. doi:10.1038/ng.2872.

Sasikumar, Arjun N., Winder B. Perez, and Terri Goss Kinzy. 2012. The Many Roles of the Eukaryotic Elongation Factor 1 Complex. Wires RNA 3 (4):543–555. doi:10.1002/wrna.1118.

Schmirler, Robert, Michael Heinzinger, and Burkhard Rost. 2024. Fine-Tuning Protein Language Models Boosts Predictions across Diverse Tasks. Nature Communications 15 (1). Nature Publishing Group:7407. doi:10.1038/s41467-024-51844-2.

Shi, Shao-Ping, Jian-Ding Qiu, Xing-Yu Sun, Sheng-Bao Suo, Shu-Yun Huang, and Ru-Ping Liang. 2012. PMeS: Prediction of Methylation Sites Based on Enhanced Feature Encoding Scheme. PLOS One 7 (6):e38772. doi:10.1371/journal.pone.0038772.

Shi, Yinan, Yanzhi Guo, Yayun Hu, and Menglong Li. 2015. Position-Specific Prediction of Methylation Sites from Sequence Conservation Based on Information Theory. Scientific Reports 5 (1):12403. doi:10.1038/srep12403.

Shrestha, Palistha, Jeevan Kandel, Hilal Tayara, and Kil To Chong. 2024. DL-SPhos: Prediction of Serine Phosphorylation Sites Using Transformer Language Model. Computers in Biology and Medicine 169 (February):107925. doi:10.1016/j.compbiomed.2024.107925.

Shukri, Ali H., Valentina Lukinović, François Charih, and Kyle K. Biggar. 2023. Unraveling the Battle for Lysine: A Review of the Competition among Post-Translational Modifications. Biochimica Et Biophysica Acta (BBA) - Gene Regulatory Mechanisms 1866 (4):194990. doi:10.1016/j.bbagrm.2023.194990.

Sondka, Zbyslaw, Nidhi Bindal Dhir, Denise Carvalho-Silva, Steven Jupe, Madhumita, Karen McLaren, Mike Starkey, Sari Ward, Jennifer Wilding, Madiha Ahmed, et al. 2024. COSMIC: A Curated Database of Somatic Variants and Clinical Data for Cancer. Nucleic Acids Research 52 (D1):D1210–D1217. doi:10.1093/nar/gkad986.

Spadaro, Austin, Alok Sharma, and Iman Dehzangi. 2024. Predicting Lysine Methylation Sites Using a Convolutional Neural Network. Methods 226 (June):127–132. doi:10.1016/j.ymeth.2024.04.007.

Straining, Rachael, and William Eighmy. 2022. Tazemetostat: EZH2 Inhibitor. Journal of the Advanced Practitioner in Oncology 13 (2):158. doi:10.6004/jadpro.2022.13.2.7.

Sun, Rongxin, Longyan Yang, Yan Wang, Yuanyuan Zhang, Jing Ke, and Dong Zhao. 2021. DNAJB11 Predicts a Poor Prognosis and Is Associated with Immune Infiltration in Thyroid Carcinoma: A Bioinformatics Analysis. Journal of International Medical Research 49 (11). SAGE Publications Ltd:03000605211053722. doi:10.1177/03000605211053722.

The UniProt Consortium. 2025. UniProt: The Universal Protein Knowledgebase in 2025. Nucleic Acids Research 53 (D1):D609–D617. doi:10.1093/nar/gkae1010.

Vanwetswinkel, Sophie, Jan Kriek, Gregers R. Andersen, Peter Güntert, Jan Dijk, Gerard W. Canters, and Gregg Siegal. 2003. Solution Structure of the 162 Residue C-terminal Domain of Human Elongation Factor 1Bγ. Journal of Biological Chemistry 278 (44):43443–43451. doi:10.1074/jbc.M306031200.

Vaswani, Ashish, Noam Shazeer, Niki Parmar, Jakob Uszkoreit, Llion Jones, Aidan N. Gomez, Lukasz Kaiser, and Illia Polosukhin. 2017. Attention Is All You Need. Preprint: arXiv. doi:10.48550/arXiv.1706.03762.

Xu, Kexin, Zhenhua Jeremy Wu, Anna C. Groner, Housheng Hansen He, Changmeng Cai, Rosina T. Lis, Xiaoqiu Wu, Edward C. Stack, Massimo Loda, Tao Liu, et al. 2012. EZH2 Oncogenic Activity in Castration-Resistant Prostate Cancer Cells Is Polycomb-independent. Science 338 (6113):1465–1469. doi:10.1126/science.1227604.

Xue, Yu, Fengfeng Zhou, Minjie Zhu, Kashif Ahmed, Guoliang Chen, and Xuebiao Yao. 2005. GPS: A Comprehensive Www Server for Phosphorylation Sites Prediction. Nucleic Acids Research 33 (suppl_2):W184–W187. doi:10.1093/nar/gki393.

Zheng, Wei, Qiqige Wuyun, Micah Cheng, Gang Hu, and Yanping Zhang. 2020. Two-Level Protein Methylation Prediction Using Structure Model-Based Features. Scientific Reports 10 (1):6008. doi:10.1038/s41598-020-62883-2.

