## Supplemental Table 1 for "Leveraging learned representations and multitask learning for lysine methylation site discovery"

Table S.1: Isolation list of 100 putative methylation sites identified with MethylSight 2.0 and selected for *in vitro* validation

| | **Protein (Uniprot ID)** | **Position** | **Score** | **Peptide** | **Modified** | **Methyl state** | **m/z (+2)** | **m/z (+3)** | **m/z (+4)** |
|---|---|---|---|---|---|---|---|---|---|
| 1 | A0A024R1R8 | 8 | 0.962 | MSSHEGGK | False | null | 416.685 | 278.125 | 208.846 |
| 2 | A0A024R1R8 | 8 | 0.962 | MSSHEGGK | False | mono | 423.692 | 282.797 | 212.35 |
| 3 | A0A024R1R8 | 8 | 0.962 | MSSHEGGK | False | di | 430.7 | 287.469 | 215.854 |
| 4 | A0A024R1R8 | 8 | 0.962 | MSSHEGGK | False | tri | 437.708 | 292.141 | 219.358 |
| 5 | A2PYH4 | 1395 | 0.914 | NPNSSNYK | False | null | 462.215 | 308.479 | 231.611 |
| 6 | A2PYH4 | 1395 | 0.914 | NPNSSNYK | False | mono | 469.222 | 313.151 | 235.115 |
| 7 | A2PYH4 | 1395 | 0.914 | NPNSSNYK | False | di | 476.23 | 317.823 | 238.619 |
| 8 | A2PYH4 | 1395 | 0.914 | NPNSSNYK | False | tri | 483.238 | 322.494 | 242.123 |
| 9 | A4D1E9 | 376 | 0.935 | QLLNLWISDTMSSTEPPSK | False | null | 1074.038 | 716.361 | 537.523 |
| 10 | A4D1E9 | 376 | 0.935 | QLLNLWISDTMSSTEPPSK | False | mono | 1081.046 | 721.033 | 541.027 |
| 11 | A4D1E9 | 376 | 0.935 | QLLNLWISDTMSSTEPPSK | False | di | 1088.054 | 725.705 | 544.531 |
| 12 | A4D1E9 | 376 | 0.935 | QLLNLWISDTMSSTEPPSK | False | tri | 1095.062 | 730.377 | 548.034 |
| 13 | B2CW77 | 75 | 0.957 | VSLVGELSK | False | null | 466.277 | 311.187 | 233.642 |
| 14 | B2CW77 | 75 | 0.957 | VSLVGELSK | False | mono | 473.284 | 315.859 | 237.146 |
| 15 | B2CW77 | 75 | 0.957 | VSLVGELSK | False | di | 480.292 | 320.531 | 240.65 |
| 16 | B2CW77 | 75 | 0.957 | VSLVGELSK | False | tri | 487.3 | 325.202 | 244.154 |
| 17 | O00300 | 255 | 0.912 | QHSSQEQTFQLLK | False | null | 787.402 | 525.27 | 394.205 |
| 18 | O00300 | 255 | 0.912 | QHSSQEQTFQLLK | False | mono | 794.41 | 529.942 | 397.709 |
| 19 | O00300 | 255 | 0.912 | QHSSQEQTFQLLK | False | di | 801.418 | 534.614 | 401.213 |
| 20 | O00300 | 255 | 0.912 | QHSSQEQTFQLLK | False | tri | 808.426 | 539.286 | 404.716 |
| 21 | O14966 | 20 | 0.923 | VLVVGDAAVGK | True | null | 514.311 | 343.21 | 257.659 |
| 22 | O14966 | 20 | 0.923 | VLVVGDAAVGK | True | mono | 521.319 | 347.882 | 261.163 |
| 23 | O14966 | 20 | 0.923 | VLVVGDAAVGK | True | di | 528.327 | 352.553 | 264.667 |
| 24 | O14966 | 20 | 0.923 | VLVVGDAAVGK | True | tri | 535.334 | 357.225 | 268.171 |
| 25 | O15144 | 256 | 0.909 | DYLHYHIK | True | null | 544.78 | 363.522 | 272.894 |
| 26 | O15144 | 256 | 0.909 | DYLHYHIK | True | mono | 551.788 | 368.194 | 276.397 |
| 27 | O15144 | 256 | 0.909 | DYLHYHIK | True | di | 558.795 | 372.866 | 279.901 |
| 28 | O15144 | 256 | 0.909 | DYLHYHIK | True | tri | 565.803 | 377.538 | 283.405 |
| 29 | O43505 | 394 | 0.942 | EAENQHNK | True | null | 485.223 | 323.818 | 243.115 |
| 30 | O43505 | 394 | 0.942 | EAENQHNK | True | mono | 492.231 | 328.49 | 246.619 |
| 31 | O43505 | 394 | 0.942 | EAENQHNK | True | di | 499.239 | 333.161 | 250.123 |
| 32 | O43505 | 394 | 0.942 | EAENQHNK | True | tri | 506.246 | 337.833 | 253.627 |
| 33 | O75129 | 1337 | 0.917 | NTYGESK | False | null | 399.685 | 266.792 | 200.346 |
| 34 | O75129 | 1337 | 0.917 | NTYGESK | False | mono | 406.693 | 271.464 | 203.85 |
| 35 | O75129 | 1337 | 0.917 | NTYGESK | False | di | 413.701 | 276.136 | 207.354 |
| 36 | O75129 | 1337 | 0.917 | NTYGESK | False | tri | 420.709 | 280.808 | 210.858 |
| 37 | O75223 | 181 | 0.976 | VSEEIEDIIK | False | null | 587.814 | 392.212 | 294.41 |
| 38 | O75223 | 181 | 0.976 | VSEEIEDIIK | False | mono | 594.822 | 396.883 | 297.914 |
| 39 | O75223 | 181 | 0.976 | VSEEIEDIIK | False | di | 601.829 | 401.555 | 301.418 |
| 40 | O75223 | 181 | 0.976 | VSEEIEDIIK | False | tri | 608.837 | 406.227 | 304.922 |

| | Protein (Uniprot ID) | Position | Score | Peptide | Modified | Methyl state | m/z (+2) | m/z (+3) | m/z (+4) |
|---|---|---|---|---|---|---|---|---|---|
| 41 | O75608 | 105 | 0.913 | ALIDQEVK | True | null | 458.261 | 305.843 | 229.634 |
| 42 | O75608 | 105 | 0.913 | ALIDQEVK | True | mono | 465.269 | 310.515 | 233.138 |
| 43 | O75608 | 105 | 0.913 | ALIDQEVK | True | di | 472.277 | 315.187 | 236.642 |
| 44 | O75608 | 105 | 0.913 | ALIDQEVK | True | tri | 479.284 | 319.859 | 240.146 |
| 45 | O75934 | 218 | 0.932 | QQHGEANK | True | null | 456.22 | 304.483 | 228.614 |
| 46 | O75934 | 218 | 0.932 | QQHGEANK | True | mono | 463.228 | 309.154 | 232.118 |
| 47 | O75934 | 218 | 0.932 | QQHGEANK | True | di | 470.236 | 313.826 | 235.622 |
| 48 | O75934 | 218 | 0.932 | QQHGEANK | True | tri | 477.244 | 318.498 | 239.125 |
| 49 | O95372 | 69 | 0.918 | IPVTLNMK | False | null | 458.27 | 305.849 | 229.639 |
| 50 | O95372 | 69 | 0.918 | IPVTLNMK | False | mono | 465.278 | 310.521 | 233.143 |
| 51 | O95372 | 69 | 0.918 | IPVTLNMK | False | di | 472.286 | 315.193 | 236.647 |
| 52 | O95372 | 69 | 0.918 | IPVTLNMK | False | tri | 479.294 | 319.865 | 240.15 |
| 53 | P02794 | 120 | 0.964 | NVNQSLLELHK | True | null | 647.859 | 432.242 | 324.433 |
| 54 | P02794 | 120 | 0.964 | NVNQSLLELHK | True | mono | 654.867 | 436.914 | 327.937 |
| 55 | P02794 | 120 | 0.964 | NVNQSLLELHK | True | di | 661.875 | 441.586 | 331.441 |
| 56 | P02794 | 120 | 0.964 | NVNQSLLELHK | True | tri | 668.883 | 446.258 | 334.945 |
| 57 | P06276 | 586 | 0.934 | WNNYMMDWK | False | null | 644.268 | 429.848 | 322.638 |
| 58 | P06276 | 586 | 0.934 | WNNYMMDWK | False | mono | 651.276 | 434.52 | 326.142 |
| 59 | P06276 | 586 | 0.934 | WNNYMMDWK | False | di | 658.284 | 439.191 | 329.645 |
| 60 | P06276 | 586 | 0.934 | WNNYMMDWK | False | tri | 665.291 | 443.863 | 333.149 |
| 61 | P08590 | 107 | 0.967 | QEELNTK | False | null | 431.219 | 287.815 | 216.113 |
| 62 | P08590 | 107 | 0.967 | QEELNTK | False | mono | 438.227 | 292.487 | 219.617 |
| 63 | P08590 | 107 | 0.967 | QEELNTK | False | di | 445.235 | 297.159 | 223.121 |
| 64 | P08590 | 107 | 0.967 | QEELNTK | False | tri | 452.243 | 301.831 | 226.625 |
| 65 | P0C0P6 | 80 | 0.987 | NGVGTGMK | False | null | 382.192 | 255.13 | 191.6 |
| 66 | P0C0P6 | 80 | 0.987 | NGVGTGMK | False | mono | 389.2 | 259.802 | 195.104 |
| 67 | P0C0P6 | 80 | 0.987 | NGVGTGMK | False | di | 396.208 | 264.474 | 198.607 |
| 68 | P0C0P6 | 80 | 0.987 | NGVGTGMK | False | tri | 403.215 | 269.146 | 202.111 |
| 69 | P0DME0 | 13 | 0.962 | QSPLPLQK | False | null | 455.771 | 304.183 | 228.389 |
| 70 | P0DME0 | 13 | 0.962 | QSPLPLQK | False | mono | 462.779 | 308.855 | 231.893 |
| 71 | P0DME0 | 13 | 0.962 | QSPLPLQK | False | di | 469.787 | 313.527 | 235.397 |
| 72 | P0DME0 | 13 | 0.962 | QSPLPLQK | False | tri | 476.795 | 318.199 | 238.901 |
| 73 | P0DP57 | 66 | 0.922 | VLSNTEDLPLVTK | False | null | 714.901 | 476.936 | 357.954 |
| 74 | P0DP57 | 66 | 0.922 | VLSNTEDLPLVTK | False | mono | 721.909 | 481.608 | 361.458 |
| 75 | P0DP57 | 66 | 0.922 | VLSNTEDLPLVTK | False | di | 728.917 | 486.28 | 364.962 |
| 76 | P0DP57 | 66 | 0.922 | VLSNTEDLPLVTK | False | tri | 735.924 | 490.952 | 368.466 |
| 77 | P11233 | 159 | 0.955 | AEQWNVNYVETSAK | True | null | 819.892 | 546.93 | 410.449 |
| 78 | P11233 | 159 | 0.955 | AEQWNVNYVETSAK | True | mono | 826.899 | 551.602 | 413.953 |
| 79 | P11233 | 159 | 0.955 | AEQWNVNYVETSAK | True | di | 833.907 | 556.274 | 417.457 |
| 80 | P11233 | 159 | 0.955 | AEQWNVNYVETSAK | True | tri | 840.915 | 560.946 | 420.961 |
| 81 | P13073 | 78 | 0.907 | ASWSSLSMDEK | False | null | 620.779 | 414.189 | 310.893 |
| 82 | P13073 | 78 | 0.907 | ASWSSLSMDEK | False | mono | 627.787 | 418.861 | 314.397 |
| 83 | P13073 | 78 | 0.907 | ASWSSLSMDEK | False | di | 634.795 | 423.532 | 317.901 |
| 84 | P13073 | 78 | 0.907 | ASWSSLSMDEK | False | tri | 641.803 | 428.204 | 321.405 |
| 85 | P20472 | 37 | 0.975 | FFQMVGLK | False | null | 485.265 | 323.846 | 243.136 |

| | Protein (Uniprot ID) | Position | Score | Peptide | Modified | Methyl state | m/z (+2) | m/z (+3) | m/z (+4) |
|---|---|---|---|---|---|---|---|---|---|
| 86 | P20472 | 37 | 0.975 | FFQMVGLK | False | mono | 492.273 | 328.518 | 246.64 |
| 87 | P20472 | 37 | 0.975 | FFQMVGLK | False | di | 499.281 | 333.189 | 250.144 |
| 88 | P20472 | 37 | 0.975 | FFQMVGLK | False | tri | 506.288 | 337.861 | 253.648 |
| 89 | P25686 | 59 | 0.934 | EVAEAYEVLSDK | True | null | 676.833 | 451.558 | 338.92 |
| 90 | P25686 | 59 | 0.934 | EVAEAYEVLSDK | True | mono | 683.84 | 456.229 | 342.424 |
| 91 | P25686 | 59 | 0.934 | EVAEAYEVLSDK | True | di | 690.848 | 460.901 | 345.928 |
| 92 | P25686 | 59 | 0.934 | EVAEAYEVLSDK | True | tri | 697.856 | 465.573 | 349.432 |
| 93 | P25705 | 539 | 0.947 | ISEQSDAK | True | null | 439.217 | 293.147 | 220.112 |
| 94 | P25705 | 539 | 0.947 | ISEQSDAK | True | mono | 446.225 | 297.819 | 223.616 |
| 95 | P25705 | 539 | 0.947 | ISEQSDAK | True | di | 453.232 | 302.491 | 227.12 |
| 96 | P25705 | 539 | 0.947 | ISEQSDAK | True | tri | 460.24 | 307.163 | 230.624 |
| 97 | P25787 | 50 | 0.957 | AANGVVLATEK | True | null | 536.803 | 358.205 | 268.905 |
| 98 | P25787 | 50 | 0.957 | AANGVVLATEK | True | mono | 543.811 | 362.877 | 272.409 |
| 99 | P25787 | 50 | 0.957 | AANGVVLATEK | True | di | 550.819 | 367.549 | 275.913 |
| 100 | P25787 | 50 | 0.957 | AANGVVLATEK | True | tri | 557.827 | 372.22 | 279.417 |
| 101 | P26447 | 18 | 0.947 | ALDVMVSTFHK | True | null | 624.326 | 416.553 | 312.667 |
| 102 | P26447 | 18 | 0.947 | ALDVMVSTFHK | True | mono | 631.334 | 421.225 | 316.171 |
| 103 | P26447 | 18 | 0.947 | ALDVMVSTFHK | True | di | 638.342 | 425.897 | 319.675 |
| 104 | P26447 | 18 | 0.947 | ALDVMVSTFHK | True | tri | 645.35 | 430.569 | 323.179 |
| 105 | P26641 | 428 | 0.936 | EYFSWEGAFQHVGK | True | null | 842.891 | 562.263 | 421.949 |
| 106 | P26641 | 428 | 0.936 | EYFSWEGAFQHVGK | True | mono | 849.899 | 566.935 | 425.453 |
| 107 | P26641 | 428 | 0.936 | EYFSWEGAFQHVGK | True | di | 856.907 | 571.607 | 428.957 |
| 108 | P26641 | 428 | 0.936 | EYFSWEGAFQHVGK | True | tri | 863.915 | 576.279 | 432.461 |
| 109 | P27216 | 284 | 0.963 | AEVDLQGIK | False | null | 486.772 | 324.85 | 243.889 |
| 110 | P27216 | 284 | 0.963 | AEVDLQGIK | False | mono | 493.779 | 329.522 | 247.393 |
| 111 | P27216 | 284 | 0.963 | AEVDLQGIK | False | di | 500.787 | 334.194 | 250.897 |
| 112 | P27216 | 284 | 0.963 | AEVDLQGIK | False | tri | 507.795 | 338.866 | 254.401 |
| 113 | P27348 | 157 | 0.97 | QTIDNSQGAYQEAFDISK | True | null | 1007.971 | 672.317 | 504.489 |
| 114 | P27348 | 157 | 0.97 | QTIDNSQGAYQEAFDISK | True | mono | 1014.979 | 676.988 | 507.993 |
| 115 | P27348 | 157 | 0.97 | QTIDNSQGAYQEAFDISK | True | di | 1021.987 | 681.66 | 511.497 |
| 116 | P27348 | 157 | 0.97 | QTIDNSQGAYQEAFDISK | True | tri | 1028.995 | 686.332 | 515.001 |
| 117 | P28340 | 1007 | 0.942 | VGGLLAFAK | True | null | 438.271 | 292.516 | 219.639 |
| 118 | P28340 | 1007 | 0.942 | VGGLLAFAK | True | mono | 445.279 | 297.188 | 223.143 |
| 119 | P28340 | 1007 | 0.942 | VGGLLAFAK | True | di | 452.287 | 301.86 | 226.647 |
| 120 | P28340 | 1007 | 0.942 | VGGLLAFAK | True | tri | 459.295 | 306.532 | 230.151 |
| 121 | P30084 | 101 | 0.972 | AFAAGADIK | True | null | 432.235 | 288.492 | 216.621 |
| 122 | P30084 | 101 | 0.972 | AFAAGADIK | True | mono | 439.243 | 293.164 | 220.125 |
| 123 | P30084 | 101 | 0.972 | AFAAGADIK | True | di | 446.25 | 297.836 | 223.629 |
| 124 | P30084 | 101 | 0.972 | AFAAGADIK | True | tri | 453.258 | 302.508 | 227.133 |
| 125 | P46734 | 340 | 0.937 | TDIAAFVK | True | null | 432.745 | 288.832 | 216.876 |
| 126 | P46734 | 340 | 0.937 | TDIAAFVK | True | mono | 439.753 | 293.504 | 220.38 |
| 127 | P46734 | 340 | 0.937 | TDIAAFVK | True | di | 446.761 | 298.176 | 223.884 |
| 128 | P46734 | 340 | 0.937 | TDIAAFVK | True | tri | 453.768 | 302.848 | 227.388 |
| 129 | P49137 | 371 | 0.947 | VDYEQIK | True | null | 447.732 | 298.824 | 224.37 |
| 130 | P49137 | 371 | 0.947 | VDYEQIK | True | mono | 454.74 | 303.496 | 227.874 |

| | Protein (Uniprot ID) | Position | Score | Peptide | Modified | Methyl state | m/z (+2) | m/z (+3) | m/z (+4) |
|---|---|---|---|---|---|---|---|---|---|
| 131 | P49137 | 371 | 0.947 | VDYEQIK | True | di | 461.748 | 308.168 | 231.377 |
| 132 | P49137 | 371 | 0.947 | VDYEQIK | True | tri | 468.755 | 312.839 | 234.881 |
| 133 | P51809 | 172 | 0.94 | TENLVDSSVTFK | True | null | 670.341 | 447.23 | 335.674 |
| 134 | P51809 | 172 | 0.94 | TENLVDSSVTFK | True | mono | 677.348 | 451.901 | 339.178 |
| 135 | P51809 | 172 | 0.94 | TENLVDSSVTFK | True | di | 684.356 | 456.573 | 342.682 |
| 136 | P51809 | 172 | 0.94 | TENLVDSSVTFK | True | tri | 691.364 | 461.245 | 346.186 |
| 137 | P56470 | 83 | 0.906 | VVFNTLQGGK | True | null | 531.801 | 354.87 | 266.404 |
| 138 | P56470 | 83 | 0.906 | VVFNTLQGGK | True | mono | 538.809 | 359.541 | 269.908 |
| 139 | P56470 | 83 | 0.906 | VVFNTLQGGK | True | di | 545.816 | 364.213 | 273.412 |
| 140 | P56470 | 83 | 0.906 | VVFNTLQGGK | True | tri | 552.824 | 368.885 | 276.916 |
| 141 | P58546 | 11 | 0.916 | EFMWALK | True | null | 462.736 | 308.826 | 231.872 |
| 142 | P58546 | 11 | 0.916 | EFMWALK | True | mono | 469.744 | 313.498 | 235.376 |
| 143 | P58546 | 11 | 0.916 | EFMWALK | True | di | 476.752 | 318.17 | 238.879 |
| 144 | P58546 | 11 | 0.916 | EFMWALK | True | tri | 483.76 | 322.842 | 242.383 |
| 145 | P59998 | 166 | 0.969 | IVAEEFLK | True | null | 474.774 | 316.852 | 237.89 |
| 146 | P59998 | 166 | 0.969 | IVAEEFLK | True | mono | 481.781 | 321.523 | 241.394 |
| 147 | P59998 | 166 | 0.969 | IVAEEFLK | True | di | 488.789 | 326.195 | 244.898 |
| 148 | P59998 | 166 | 0.969 | IVAEEFLK | True | tri | 495.797 | 330.867 | 248.402 |
| 149 | P60228 | 407 | 0.906 | LGHVVMGNNAVSPYQQVIEK | True | null | 1092.068 | 728.381 | 546.537 |
| 150 | P60228 | 407 | 0.906 | LGHVVMGNNAVSPYQQVIEK | True | mono | 1099.075 | 733.053 | 550.041 |
| 151 | P60228 | 407 | 0.906 | LGHVVMGNNAVSPYQQVIEK | True | di | 1106.083 | 737.725 | 553.545 |
| 152 | P60228 | 407 | 0.906 | LGHVVMGNNAVSPYQQVIEK | True | tri | 1113.091 | 742.396 | 557.049 |
| 153 | P61244 | 24 | 0.956 | FQSAADK | True | null | 383.69 | 256.129 | 192.349 |
| 154 | P61244 | 24 | 0.956 | FQSAADK | True | mono | 390.698 | 260.801 | 195.853 |
| 155 | P61244 | 24 | 0.956 | FQSAADK | True | di | 397.706 | 265.473 | 199.357 |
| 156 | P61244 | 24 | 0.956 | FQSAADK | True | tri | 404.714 | 270.145 | 202.86 |
| 157 | P61758 | 59 | 0.971 | LDEQYQK | True | null | 462.227 | 308.487 | 231.617 |
| 158 | P61758 | 59 | 0.971 | LDEQYQK | True | mono | 469.235 | 313.159 | 235.121 |
| 159 | P61758 | 59 | 0.971 | LDEQYQK | True | di | 476.243 | 317.831 | 238.625 |
| 160 | P61758 | 59 | 0.971 | LDEQYQK | True | tri | 483.251 | 322.503 | 242.129 |
| 161 | P98175 | 915 | 0.942 | GSSYGVTSTESYK | False | null | 683.312 | 455.877 | 342.16 |
| 162 | P98175 | 915 | 0.942 | GSSYGVTSTESYK | False | mono | 690.32 | 460.549 | 345.664 |
| 163 | P98175 | 915 | 0.942 | GSSYGVTSTESYK | False | di | 697.328 | 465.221 | 349.167 |
| 164 | P98175 | 915 | 0.942 | GSSYGVTSTESYK | False | tri | 704.336 | 469.893 | 352.671 |
| 165 | Q08623 | 123 | 0.93 | HGIPFALATSSGSASFDMK | False | null | 962.467 | 641.98 | 481.737 |
| 166 | Q08623 | 123 | 0.93 | HGIPFALATSSGSASFDMK | False | mono | 969.475 | 646.652 | 485.241 |
| 167 | Q08623 | 123 | 0.93 | HGIPFALATSSGSASFDMK | False | di | 976.483 | 651.324 | 488.745 |
| 168 | Q08623 | 123 | 0.93 | HGIPFALATSSGSASFDMK | False | tri | 983.491 | 655.996 | 492.249 |
| 169 | Q0VDE8 | 75 | 0.944 | GTLHGQEK | False | null | 435.227 | 290.487 | 218.117 |
| 170 | Q0VDE8 | 75 | 0.944 | GTLHGQEK | False | mono | 442.235 | 295.159 | 221.621 |
| 171 | Q0VDE8 | 75 | 0.944 | GTLHGQEK | False | di | 449.243 | 299.831 | 225.125 |
| 172 | Q0VDE8 | 75 | 0.944 | GTLHGQEK | False | tri | 456.251 | 304.503 | 228.629 |
| 173 | Q13043 | 480 | 0.964 | QPILDAIEAK | False | null | 549.314 | 366.545 | 275.16 |
| 174 | Q13043 | 480 | 0.964 | QPILDAIEAK | False | mono | 556.322 | 371.217 | 278.664 |
| 175 | Q13043 | 480 | 0.964 | QPILDAIEAK | False | di | 563.329 | 375.889 | 282.168 |

| | Protein (Uniprot ID) | Position | Score | Peptide | Modified | Methyl state | m/z (+2) | m/z (+3) | m/z (+4) |
|---|---|---|---|---|---|---|---|---|---|
| 176 | Q13043 | 480 | 0.964 | QPILDAIEAK | False | tri | 570.337 | 380.561 | 285.672 |
| 177 | Q13823 | 721 | 0.933 | TNDSEGQK | False | null | 439.696 | 293.467 | 220.352 |
| 178 | Q13823 | 721 | 0.933 | TNDSEGQK | False | mono | 446.704 | 298.138 | 223.856 |
| 179 | Q13823 | 721 | 0.933 | TNDSEGQK | False | di | 453.712 | 302.81 | 227.36 |
| 180 | Q13823 | 721 | 0.933 | TNDSEGQK | False | tri | 460.72 | 307.482 | 230.863 |
| 181 | Q14914 | 318 | 0.955 | EYIIEGFENMPAAFMGMLK | False | null | 1096.017 | 731.014 | 548.512 |
| 182 | Q14914 | 318 | 0.955 | EYIIEGFENMPAAFMGMLK | False | mono | 1103.025 | 735.686 | 552.016 |
| 183 | Q14914 | 318 | 0.955 | EYIIEGFENMPAAFMGMLK | False | di | 1110.033 | 740.358 | 555.52 |
| 184 | Q14914 | 318 | 0.955 | EYIIEGFENMPAAFMGMLK | False | tri | 1117.041 | 745.03 | 559.024 |
| 185 | Q15286 | 189 | 0.969 | QQQQQQNDVVK | True | null | 671.839 | 448.229 | 336.423 |
| 186 | Q15286 | 189 | 0.969 | QQQQQQNDVVK | True | mono | 678.847 | 452.9 | 339.927 |
| 187 | Q15286 | 189 | 0.969 | QQQQQQNDVVK | True | di | 685.855 | 457.572 | 343.431 |
| 188 | Q15286 | 189 | 0.969 | QQQQQQNDVVK | True | tri | 692.863 | 462.244 | 346.935 |
| 189 | Q15506 | 49 | 0.897 | EQPDNIPAFAAAYFESLLEK | False | null | 1127.057 | 751.707 | 564.032 |
| 190 | Q15506 | 49 | 0.897 | EQPDNIPAFAAAYFESLLEK | False | mono | 1134.065 | 756.379 | 567.536 |
| 191 | Q15506 | 49 | 0.897 | EQPDNIPAFAAAYFESLLEK | False | di | 1141.073 | 761.051 | 571.04 |
| 192 | Q15506 | 49 | 0.897 | EQPDNIPAFAAAYFESLLEK | False | tri | 1148.081 | 765.723 | 574.544 |
| 193 | Q16836 | 249 | 0.9 | EDIDTAMK | True | null | 461.713 | 308.144 | 231.36 |
| 194 | Q16836 | 249 | 0.9 | EDIDTAMK | True | mono | 468.721 | 312.816 | 234.864 |
| 195 | Q16836 | 249 | 0.9 | EDIDTAMK | True | di | 475.729 | 317.488 | 238.368 |
| 196 | Q16836 | 249 | 0.9 | EDIDTAMK | True | tri | 482.736 | 322.16 | 241.872 |
| 197 | Q32NC0 | 85 | 0.96 | NYTLSFK | False | null | 436.729 | 291.489 | 218.868 |
| 198 | Q32NC0 | 85 | 0.96 | NYTLSFK | False | mono | 443.737 | 296.16 | 222.372 |
| 199 | Q32NC0 | 85 | 0.96 | NYTLSFK | False | di | 450.745 | 300.832 | 225.876 |
| 200 | Q32NC0 | 85 | 0.96 | NYTLSFK | False | tri | 457.753 | 305.504 | 229.38 |
| 201 | Q5MAI5 | 33 | 0.93 | TSGQVVAVK | True | null | 444.761 | 296.843 | 222.884 |
| 202 | Q5MAI5 | 33 | 0.93 | TSGQVVAVK | True | mono | 451.769 | 301.515 | 226.388 |
| 203 | Q5MAI5 | 33 | 0.93 | TSGQVVAVK | True | di | 458.777 | 306.187 | 229.892 |
| 204 | Q5MAI5 | 33 | 0.93 | TSGQVVAVK | True | tri | 465.785 | 310.859 | 233.396 |
| 205 | Q5TC84 | 344 | 0.918 | MSSPLASSHNSQTSMHK | False | null | 915.417 | 610.614 | 458.212 |
| 206 | Q5TC84 | 344 | 0.918 | MSSPLASSHNSQTSMHK | False | mono | 922.425 | 615.286 | 461.716 |
| 207 | Q5TC84 | 344 | 0.918 | MSSPLASSHNSQTSMHK | False | di | 929.433 | 619.958 | 465.22 |
| 208 | Q5TC84 | 344 | 0.918 | MSSPLASSHNSQTSMHK | False | tri | 936.441 | 624.629 | 468.724 |
| 209 | Q5VT25 | 1647 | 0.954 | SSAQNGSALK | False | null | 481.749 | 321.502 | 241.378 |
| 210 | Q5VT25 | 1647 | 0.954 | SSAQNGSALK | False | mono | 488.757 | 326.173 | 244.882 |
| 211 | Q5VT25 | 1647 | 0.954 | SSAQNGSALK | False | di | 495.764 | 330.845 | 248.386 |
| 212 | Q5VT25 | 1647 | 0.954 | SSAQNGSALK | False | tri | 502.772 | 335.517 | 251.89 |
| 213 | Q6DD88 | 399 | 0.958 | QLALDHFK | False | null | 486.269 | 324.515 | 243.638 |
| 214 | Q6DD88 | 399 | 0.958 | QLALDHFK | False | mono | 493.277 | 329.187 | 247.142 |
| 215 | Q6DD88 | 399 | 0.958 | QLALDHFK | False | di | 500.285 | 333.859 | 250.646 |
| 216 | Q6DD88 | 399 | 0.958 | QLALDHFK | False | tri | 507.293 | 338.531 | 254.15 |
| 217 | Q6IQ20 | 18 | 0.906 | MDENESNQSLMTSSQYPK | False | null | 1044.946 | 696.966 | 522.977 |
| 218 | Q6IQ20 | 18 | 0.906 | MDENESNQSLMTSSQYPK | False | mono | 1051.954 | 701.638 | 526.481 |
| 219 | Q6IQ20 | 18 | 0.906 | MDENESNQSLMTSSQYPK | False | di | 1058.962 | 706.31 | 529.984 |
| 220 | Q6IQ20 | 18 | 0.906 | MDENESNQSLMTSSQYPK | False | tri | 1065.97 | 710.982 | 533.488 |

|  | Protein (Uniprot ID) | Position | Score | Peptide | Modified | Methyl state | m/z (+2) | m/z (+3) | m/z (+4) |
|---|---|---|---|---|---|---|---|---|---|
| 221 | Q6P2I3 | 18 | 0.929 | LLTALLQAQK | False | null | 549.848 | 366.901 | 275.427 |
| 222 | Q6P2I3 | 18 | 0.929 | LLTALLQAQK | False | mono | 556.856 | 371.573 | 278.931 |
| 223 | Q6P2I3 | 18 | 0.929 | LLTALLQAQK | False | di | 563.863 | 376.245 | 282.435 |
| 224 | Q6P2I3 | 18 | 0.929 | LLTALLQAQK | False | tri | 570.871 | 380.917 | 285.939 |
| 225 | Q7Z6K5 | 203 | 0.958 | TGASWTDNIMAQK | True | null | 711.838 | 474.894 | 356.423 |
| 226 | Q7Z6K5 | 203 | 0.958 | TGASWTDNIMAQK | True | mono | 718.846 | 479.566 | 359.926 |
| 227 | Q7Z6K5 | 203 | 0.958 | TGASWTDNIMAQK | True | di | 725.853 | 484.238 | 363.43 |
| 228 | Q7Z6K5 | 203 | 0.958 | TGASWTDNIMAQK | True | tri | 732.861 | 488.91 | 366.934 |
| 229 | Q86SX6 | 151 | 0.93 | LGIHSALLDEK | False | null | 598.338 | 399.228 | 299.672 |
| 230 | Q86SX6 | 151 | 0.93 | LGIHSALLDEK | False | mono | 605.346 | 403.899 | 303.176 |
| 231 | Q86SX6 | 151 | 0.93 | LGIHSALLDEK | False | di | 612.353 | 408.571 | 306.68 |
| 232 | Q86SX6 | 151 | 0.93 | LGIHSALLDEK | False | tri | 619.361 | 413.243 | 310.184 |
| 233 | Q8IX90 | 389 | 0.954 | YNSNLATPIAIK | False | null | 652.864 | 435.578 | 326.936 |
| 234 | Q8IX90 | 389 | 0.954 | YNSNLATPIAIK | False | mono | 659.872 | 440.25 | 330.44 |
| 235 | Q8IX90 | 389 | 0.954 | YNSNLATPIAIK | False | di | 666.88 | 444.922 | 333.944 |
| 236 | Q8IX90 | 389 | 0.954 | YNSNLATPIAIK | False | tri | 673.888 | 449.594 | 337.447 |
| 237 | Q8IY31 | 34 | 0.923 | VLDPEVTQQTIELK | True | null | 806.943 | 538.298 | 403.975 |
| 238 | Q8IY31 | 34 | 0.923 | VLDPEVTQQTIELK | True | mono | 813.951 | 542.97 | 407.479 |
| 239 | Q8IY31 | 34 | 0.923 | VLDPEVTQQTIELK | True | di | 820.959 | 547.642 | 410.983 |
| 240 | Q8IY31 | 34 | 0.923 | VLDPEVTQQTIELK | True | tri | 827.967 | 552.314 | 414.487 |
| 241 | Q8N118 | 504 | 0.938 | NGMYLHLK | False | null | 488.258 | 325.841 | 244.632 |
| 242 | Q8N118 | 504 | 0.938 | NGMYLHLK | False | mono | 495.265 | 330.513 | 248.136 |
| 243 | Q8N118 | 504 | 0.938 | NGMYLHLK | False | di | 502.273 | 335.185 | 251.64 |
| 244 | Q8N118 | 504 | 0.938 | NGMYLHLK | False | tri | 509.281 | 339.857 | 255.144 |
| 245 | Q8N4H5 | 46 | 0.979 | VTPFILK | True | null | 409.263 | 273.178 | 205.135 |
| 246 | Q8N4H5 | 46 | 0.979 | VTPFILK | True | mono | 416.271 | 277.849 | 208.639 |
| 247 | Q8N4H5 | 46 | 0.979 | VTPFILK | True | di | 423.278 | 282.521 | 212.143 |
| 248 | Q8N4H5 | 46 | 0.979 | VTPFILK | True | tri | 430.286 | 287.193 | 215.647 |
| 249 | Q8N8Q3 | 170 | 0.905 | LLQVDGLENNALHK | True | null | 782.428 | 521.954 | 391.718 |
| 250 | Q8N8Q3 | 170 | 0.905 | LLQVDGLENNALHK | True | mono | 789.436 | 526.626 | 395.222 |
| 251 | Q8N8Q3 | 170 | 0.905 | LLQVDGLENNALHK | True | di | 796.444 | 531.298 | 398.726 |
| 252 | Q8N8Q3 | 170 | 0.905 | LLQVDGLENNALHK | True | tri | 803.452 | 535.97 | 402.229 |
| 253 | Q8NA97 | 84 | 0.91 | LFIYSSK | False | null | 429.242 | 286.497 | 215.125 |
| 254 | Q8NA97 | 84 | 0.91 | LFIYSSK | False | mono | 436.25 | 291.169 | 218.629 |
| 255 | Q8NA97 | 84 | 0.91 | LFIYSSK | False | di | 443.258 | 295.841 | 222.132 |
| 256 | Q8NA97 | 84 | 0.91 | LFIYSSK | False | tri | 450.265 | 300.513 | 225.636 |
| 257 | Q8NE09 | 1254 | 0.942 | ASSSTMSLK | False | null | 456.229 | 304.488 | 228.618 |
| 258 | Q8NE09 | 1254 | 0.942 | ASSSTMSLK | False | mono | 463.237 | 309.16 | 232.122 |
| 259 | Q8NE09 | 1254 | 0.942 | ASSSTMSLK | False | di | 470.244 | 313.832 | 235.626 |
| 260 | Q8NE09 | 1254 | 0.942 | ASSSTMSLK | False | tri | 477.252 | 318.504 | 239.13 |
| 261 | Q8NEF9 | 378 | 0.942 | SLDFPQNEPQIK | True | null | 708.362 | 472.577 | 354.685 |
| 262 | Q8NEF9 | 378 | 0.942 | SLDFPQNEPQIK | True | mono | 715.37 | 477.249 | 358.189 |
| 263 | Q8NEF9 | 378 | 0.942 | SLDFPQNEPQIK | True | di | 722.378 | 481.921 | 361.692 |
| 264 | Q8NEF9 | 378 | 0.942 | SLDFPQNEPQIK | True | tri | 729.385 | 486.593 | 365.196 |
| 265 | Q8NFR3 | 66 | 0.909 | LAWEFFSK | False | null | 514.266 | 343.18 | 257.637 |

| | Protein (Uniprot ID) | Position | Score | Peptide | Modified | Methyl state | m/z (+2) | m/z (+3) | m/z (+4) |
|---|---|---|---|---|---|---|---|---|---|
| 266 | Q8NFR3 | 66 | 0.909 | LAWEFFSK | False | mono | 521.274 | 347.852 | 261.141 |
| 267 | Q8NFR3 | 66 | 0.909 | LAWEFFSK | False | di | 528.282 | 352.524 | 264.644 |
| 268 | Q8NFR3 | 66 | 0.909 | LAWEFFSK | False | tri | 535.289 | 357.195 | 268.148 |
| 269 | Q8NGY0 | 325 | 0.975 | MMGNTVALK | False | null | 482.751 | 322.17 | 241.879 |
| 270 | Q8NGY0 | 325 | 0.975 | MMGNTVALK | False | mono | 489.759 | 326.842 | 245.383 |
| 271 | Q8NGY0 | 325 | 0.975 | MMGNTVALK | False | di | 496.767 | 331.514 | 248.887 |
| 272 | Q8NGY0 | 325 | 0.975 | MMGNTVALK | False | tri | 503.775 | 336.186 | 252.391 |
| 273 | Q8WUD6 | 383 | 0.926 | HLHLNIFK | False | null | 511.301 | 341.203 | 256.154 |
| 274 | Q8WUD6 | 383 | 0.926 | HLHLNIFK | False | mono | 518.309 | 345.875 | 259.658 |
| 275 | Q8WUD6 | 383 | 0.926 | HLHLNIFK | False | di | 525.316 | 350.547 | 263.162 |
| 276 | Q8WUD6 | 383 | 0.926 | HLHLNIFK | False | tri | 532.324 | 355.219 | 266.666 |
| 277 | Q92499 | 702 | 0.952 | AAGGGSYK | True | null | 355.677 | 237.454 | 178.342 |
| 278 | Q92499 | 702 | 0.952 | AAGGGSYK | True | mono | 362.685 | 242.126 | 181.846 |
| 279 | Q92499 | 702 | 0.952 | AAGGGSYK | True | di | 369.693 | 246.798 | 185.35 |
| 280 | Q92499 | 702 | 0.952 | AAGGGSYK | True | tri | 376.701 | 251.469 | 188.854 |
| 281 | Q92901 | 297 | 0.952 | GPHMEDGK | False | null | 435.692 | 290.797 | 218.35 |
| 282 | Q92901 | 297 | 0.952 | GPHMEDGK | False | mono | 442.7 | 295.469 | 221.854 |
| 283 | Q92901 | 297 | 0.952 | GPHMEDGK | False | di | 449.708 | 300.141 | 225.358 |
| 284 | Q92901 | 297 | 0.952 | GPHMEDGK | False | tri | 456.716 | 304.813 | 228.862 |
| 285 | Q96CQ1 | 174 | 0.924 | VYQTDGLK | False | null | 462.245 | 308.499 | 231.626 |
| 286 | Q96CQ1 | 174 | 0.924 | VYQTDGLK | False | mono | 469.253 | 313.171 | 235.13 |
| 287 | Q96CQ1 | 174 | 0.924 | VYQTDGLK | False | di | 476.261 | 317.843 | 238.634 |
| 288 | Q96CQ1 | 174 | 0.924 | VYQTDGLK | False | tri | 483.269 | 322.515 | 242.138 |
| 289 | Q96IG2 | 408 | 0.897 | THLPNIK | False | null | 411.745 | 274.833 | 206.376 |
| 290 | Q96IG2 | 408 | 0.897 | THLPNIK | False | mono | 418.753 | 279.504 | 209.88 |
| 291 | Q96IG2 | 408 | 0.897 | THLPNIK | False | di | 425.761 | 284.176 | 213.384 |
| 292 | Q96IG2 | 408 | 0.897 | THLPNIK | False | tri | 432.769 | 288.848 | 216.888 |
| 293 | Q96IM9 | 158 | 0.925 | MPQEINYK | False | null | 511.752 | 341.504 | 256.38 |
| 294 | Q96IM9 | 158 | 0.925 | MPQEINYK | False | mono | 518.76 | 346.176 | 259.884 |
| 295 | Q96IM9 | 158 | 0.925 | MPQEINYK | False | di | 525.768 | 350.848 | 263.388 |
| 296 | Q96IM9 | 158 | 0.925 | MPQEINYK | False | tri | 532.776 | 355.52 | 266.892 |
| 297 | Q96KB5 | 8 | 0.984 | MEGISNFK | True | null | 463.226 | 309.153 | 232.117 |
| 298 | Q96KB5 | 8 | 0.984 | MEGISNFK | True | mono | 470.234 | 313.825 | 235.621 |
| 299 | Q96KB5 | 8 | 0.984 | MEGISNFK | True | di | 477.242 | 318.497 | 239.124 |
| 300 | Q96KB5 | 8 | 0.984 | MEGISNFK | True | tri | 484.25 | 323.169 | 242.628 |
| 301 | Q96PM5 | 239 | 0.934 | STVQFHILGMK | False | null | 630.842 | 420.897 | 315.925 |
| 302 | Q96PM5 | 239 | 0.934 | STVQFHILGMK | False | mono | 637.85 | 425.569 | 319.429 |
| 303 | Q96PM5 | 239 | 0.934 | STVQFHILGMK | False | di | 644.858 | 430.241 | 322.933 |
| 304 | Q96PM5 | 239 | 0.934 | STVQFHILGMK | False | tri | 651.866 | 434.913 | 326.436 |
| 305 | Q96QD9 | 261 | 0.976 | TAVPSFLTK | True | null | 482.279 | 321.855 | 241.643 |
| 306 | Q96QD9 | 261 | 0.976 | TAVPSFLTK | True | mono | 489.287 | 326.527 | 245.147 |
| 307 | Q96QD9 | 261 | 0.976 | TAVPSFLTK | True | di | 496.295 | 331.199 | 248.651 |
| 308 | Q96QD9 | 261 | 0.976 | TAVPSFLTK | True | tri | 503.303 | 335.871 | 252.155 |
| 309 | Q99757 | 147 | 0.93 | NGDVVDK | False | null | 373.688 | 249.461 | 187.347 |
| 310 | Q99757 | 147 | 0.93 | NGDVVDK | False | mono | 380.695 | 254.133 | 190.851 |

| | Protein (Uniprot ID) | Position | Score | Peptide | Modified | Methyl state | m/z (+2) | m/z (+3) | m/z (+4) |
|---|---|---|---|---|---|---|---|---|---|
| 311 | Q99757 | 147 | 0.93 | NGDVVDK | False | di | 387.703 | 258.805 | 194.355 |
| 312 | Q99757 | 147 | 0.93 | NGDVVDK | False | tri | 394.711 | 263.476 | 197.859 |
| 313 | Q9BSD7 | 73 | 0.923 | VGLEPPPGK | True | null | 447.258 | 298.508 | 224.133 |
| 314 | Q9BSD7 | 73 | 0.923 | VGLEPPPGK | True | mono | 454.266 | 303.18 | 227.637 |
| 315 | Q9BSD7 | 73 | 0.923 | VGLEPPPGK | True | di | 461.274 | 307.852 | 231.141 |
| 316 | Q9BSD7 | 73 | 0.923 | VGLEPPPGK | True | tri | 468.282 | 312.524 | 234.644 |
| 317 | Q9BTT4 | 82 | 0.915 | NPQLYTK | True | null | 432.235 | 288.492 | 216.621 |
| 318 | Q9BTT4 | 82 | 0.915 | NPQLYTK | True | mono | 439.243 | 293.164 | 220.125 |
| 319 | Q9BTT4 | 82 | 0.915 | NPQLYTK | True | di | 446.25 | 297.836 | 223.629 |
| 320 | Q9BTT4 | 82 | 0.915 | NPQLYTK | True | tri | 453.258 | 302.508 | 227.133 |
| 321 | Q9BV29 | 98 | 0.913 | GLNQEVTSK | True | null | 488.259 | 325.842 | 244.633 |
| 322 | Q9BV29 | 98 | 0.913 | GLNQEVTSK | True | mono | 495.267 | 330.514 | 248.137 |
| 323 | Q9BV29 | 98 | 0.913 | GLNQEVTSK | True | di | 502.275 | 335.185 | 251.641 |
| 324 | Q9BV29 | 98 | 0.913 | GLNQEVTSK | True | tri | 509.282 | 339.857 | 255.145 |
| 325 | Q9BVG4 | 63 | 0.939 | LISSVDPQFLK | False | null | 623.856 | 416.24 | 312.431 |
| 326 | Q9BVG4 | 63 | 0.939 | LISSVDPQFLK | False | mono | 630.864 | 420.911 | 315.935 |
| 327 | Q9BVG4 | 63 | 0.939 | LISSVDPQFLK | False | di | 637.871 | 425.583 | 319.439 |
| 328 | Q9BVG4 | 63 | 0.939 | LISSVDPQFLK | False | tri | 644.879 | 430.255 | 322.943 |
| 329 | Q9BZE2 | 407 | 0.961 | QTSAFVEGVK | False | null | 533.282 | 355.857 | 267.145 |
| 330 | Q9BZE2 | 407 | 0.961 | QTSAFVEGVK | False | mono | 540.29 | 360.529 | 270.649 |
| 331 | Q9BZE2 | 407 | 0.961 | QTSAFVEGVK | False | di | 547.298 | 365.201 | 274.153 |
| 332 | Q9BZE2 | 407 | 0.961 | QTSAFVEGVK | False | tri | 554.306 | 369.873 | 277.657 |
| 333 | Q9H3Q1 | 114 | 0.973 | DSALFVK | True | null | 390.219 | 260.481 | 195.613 |
| 334 | Q9H3Q1 | 114 | 0.973 | DSALFVK | True | mono | 397.226 | 265.153 | 199.117 |
| 335 | Q9H3Q1 | 114 | 0.973 | DSALFVK | True | di | 404.234 | 269.825 | 202.621 |
| 336 | Q9H3Q1 | 114 | 0.973 | DSALFVK | True | tri | 411.242 | 274.497 | 206.125 |
| 337 | Q9H7S9 | 141 | 0.897 | SAPGAASAAAALK | True | null | 543.301 | 362.537 | 272.154 |
| 338 | Q9H7S9 | 141 | 0.897 | SAPGAASAAAALK | True | mono | 550.309 | 367.208 | 275.658 |
| 339 | Q9H7S9 | 141 | 0.897 | SAPGAASAAAALK | True | di | 557.317 | 371.88 | 279.162 |
| 340 | Q9H7S9 | 141 | 0.897 | SAPGAASAAAALK | True | tri | 564.325 | 376.552 | 282.666 |
| 341 | Q9HAV7 | 138 | 0.923 | DDNPHLK | False | null | 419.706 | 280.14 | 210.357 |
| 342 | Q9HAV7 | 138 | 0.923 | DDNPHLK | False | mono | 426.714 | 284.812 | 213.861 |
| 343 | Q9HAV7 | 138 | 0.923 | DDNPHLK | False | di | 433.722 | 289.484 | 217.365 |
| 344 | Q9HAV7 | 138 | 0.923 | DDNPHLK | False | tri | 440.73 | 294.156 | 220.869 |
| 345 | Q9NS73 | 208 | 0.965 | TDAIFTPYPGFK | True | null | 678.845 | 452.899 | 339.926 |
| 346 | Q9NS73 | 208 | 0.965 | TDAIFTPYPGFK | True | mono | 685.853 | 457.571 | 343.43 |
| 347 | Q9NS73 | 208 | 0.965 | TDAIFTPYPGFK | True | di | 692.861 | 462.243 | 346.934 |
| 348 | Q9NS73 | 208 | 0.965 | TDAIFTPYPGFK | True | tri | 699.869 | 466.915 | 350.438 |
| 349 | Q9NSI8 | 68 | 0.915 | TSNNGGGLGK | True | null | 452.728 | 302.154 | 226.868 |
| 350 | Q9NSI8 | 68 | 0.915 | TSNNGGGLGK | True | mono | 459.736 | 306.826 | 230.371 |
| 351 | Q9NSI8 | 68 | 0.915 | TSNNGGGLGK | True | di | 466.743 | 311.498 | 233.875 |
| 352 | Q9NSI8 | 68 | 0.915 | TSNNGGGLGK | True | tri | 473.751 | 316.17 | 237.379 |
| 353 | Q9NUV7 | 530 | 0.908 | EMLDTVLEALDEMGDLLQLK | False | null | 1138.574 | 759.385 | 569.79 |
| 354 | Q9NUV7 | 530 | 0.908 | EMLDTVLEALDEMGDLLQLK | False | mono | 1145.581 | 764.057 | 573.294 |
| 355 | Q9NUV7 | 530 | 0.908 | EMLDTVLEALDEMGDLLQLK | False | di | 1152.589 | 768.729 | 576.798 |

| | Protein (Uniprot ID) | Position | Score | Peptide | Modified | Methyl state | m/z (+2) | m/z (+3) | m/z (+4) |
|---|---|---|---|---|---|---|---|---|---|
| 356 | Q9NUV7 | 530 | 0.908 | EMLDTVLEALDEMGDLLQLK | False | tri | 1159.597 | 773.401 | 580.302 |
| 357 | Q9NV56 | 200 | 0.977 | VLTANSNPSSPSAAK | True | null | 722.376 | 481.919 | 361.691 |
| 358 | Q9NV56 | 200 | 0.977 | VLTANSNPSSPSAAK | True | mono | 729.383 | 486.591 | 365.195 |
| 359 | Q9NV56 | 200 | 0.977 | VLTANSNPSSPSAAK | True | di | 736.391 | 491.263 | 368.699 |
| 360 | Q9NV56 | 200 | 0.977 | VLTANSNPSSPSAAK | True | tri | 743.399 | 495.935 | 372.203 |
| 361 | Q9NWQ9 | 128 | 0.916 | QLEFSEPDFVAK | True | null | 705.351 | 470.57 | 353.179 |
| 362 | Q9NWQ9 | 128 | 0.916 | QLEFSEPDFVAK | True | mono | 712.359 | 475.242 | 356.683 |
| 363 | Q9NWQ9 | 128 | 0.916 | QLEFSEPDFVAK | True | di | 719.367 | 479.914 | 360.187 |
| 364 | Q9NWQ9 | 128 | 0.916 | QLEFSEPDFVAK | True | tri | 726.374 | 484.585 | 363.691 |
| 365 | Q9NX63 | 173 | 0.924 | AAEEVEAK | True | null | 423.714 | 282.812 | 212.361 |
| 366 | Q9NX63 | 173 | 0.924 | AAEEVEAK | True | mono | 430.722 | 287.484 | 215.864 |
| 367 | Q9NX63 | 173 | 0.924 | AAEEVEAK | True | di | 437.729 | 292.155 | 219.368 |
| 368 | Q9NX63 | 173 | 0.924 | AAEEVEAK | True | tri | 444.737 | 296.827 | 222.872 |
| 369 | Q9NXW2 | 177 | 0.913 | QYDQFGDDK | True | null | 558.236 | 372.493 | 279.621 |
| 370 | Q9NXW2 | 177 | 0.913 | QYDQFGDDK | True | mono | 565.243 | 377.165 | 283.125 |
| 371 | Q9NXW2 | 177 | 0.913 | QYDQFGDDK | True | di | 572.251 | 381.837 | 286.629 |
| 372 | Q9NXW2 | 177 | 0.913 | QYDQFGDDK | True | tri | 579.259 | 386.508 | 290.133 |
| 373 | Q9P0L0 | 24 | 0.917 | HEQILVLDPPTDLK | True | null | 809.446 | 539.966 | 405.227 |
| 374 | Q9P0L0 | 24 | 0.917 | HEQILVLDPPTDLK | True | mono | 816.454 | 544.638 | 408.731 |
| 375 | Q9P0L0 | 24 | 0.917 | HEQILVLDPPTDLK | True | di | 823.462 | 549.31 | 412.234 |
| 376 | Q9P0L0 | 24 | 0.917 | HEQILVLDPPTDLK | True | tri | 830.469 | 553.982 | 415.738 |
| 377 | Q9UBS4 | 66 | 0.931 | NPDDPQAQEK | True | null | 571.26 | 381.176 | 286.133 |
| 378 | Q9UBS4 | 66 | 0.931 | NPDDPQAQEK | True | mono | 578.267 | 385.847 | 289.637 |
| 379 | Q9UBS4 | 66 | 0.931 | NPDDPQAQEK | True | di | 585.275 | 390.519 | 293.141 |
| 380 | Q9UBS4 | 66 | 0.931 | NPDDPQAQEK | True | tri | 592.283 | 395.191 | 296.645 |
| 381 | Q9UF47 | 184 | 0.933 | DVDFPVFLQPTNANEK | False | null | 917.455 | 611.972 | 459.231 |
| 382 | Q9UF47 | 184 | 0.933 | DVDFPVFLQPTNANEK | False | mono | 924.462 | 616.644 | 462.735 |
| 383 | Q9UF47 | 184 | 0.933 | DVDFPVFLQPTNANEK | False | di | 931.47 | 621.316 | 466.239 |
| 384 | Q9UF47 | 184 | 0.933 | DVDFPVFLQPTNANEK | False | tri | 938.478 | 625.988 | 469.743 |
| 385 | Q9UHD4 | 213 | 0.946 | HAVEGAEQWQQK | False | null | 705.842 | 470.897 | 353.424 |
| 386 | Q9UHD4 | 213 | 0.946 | HAVEGAEQWQQK | False | mono | 712.849 | 475.569 | 356.928 |
| 387 | Q9UHD4 | 213 | 0.946 | HAVEGAEQWQQK | False | di | 719.857 | 480.241 | 360.432 |
| 388 | Q9UHD4 | 213 | 0.946 | HAVEGAEQWQQK | False | tri | 726.865 | 484.913 | 363.936 |
| 389 | Q9ULV4 | 19 | 0.931 | HVFGQAVK | True | null | 443.251 | 295.836 | 222.129 |
| 390 | Q9ULV4 | 19 | 0.931 | HVFGQAVK | True | mono | 450.259 | 300.508 | 225.633 |
| 391 | Q9ULV4 | 19 | 0.931 | HVFGQAVK | True | di | 457.266 | 305.18 | 229.137 |
| 392 | Q9ULV4 | 19 | 0.931 | HVFGQAVK | True | tri | 464.274 | 309.852 | 232.641 |
| 393 | Q9Y3V2 | 267 | 0.899 | LFSEFVLALVK | False | null | 633.379 | 422.588 | 317.193 |
| 394 | Q9Y3V2 | 267 | 0.899 | LFSEFVLALVK | False | mono | 640.387 | 427.26 | 320.697 |
| 395 | Q9Y3V2 | 267 | 0.899 | LFSEFVLALVK | False | di | 647.394 | 431.932 | 324.201 |
| 396 | Q9Y3V2 | 267 | 0.899 | LFSEFVLALVK | False | tri | 654.402 | 436.604 | 327.705 |
| 397 | Q9Y6G3 | 114 | 0.935 | VEHLEEGPMIEQLSK | True | null | 869.938 | 580.294 | 435.472 |
| 398 | Q9Y6G3 | 114 | 0.935 | VEHLEEGPMIEQLSK | True | mono | 876.945 | 584.966 | 438.976 |
| 399 | Q9Y6G3 | 114 | 0.935 | VEHLEEGPMIEQLSK | True | di | 883.953 | 589.638 | 442.48 |
| 400 | Q9Y6G3 | 114 | 0.935 | VEHLEEGPMIEQLSK | True | tri | 890.961 | 594.31 | 445.984 |
