## Supplemental Table 2 for "Leveraging learned representations and multitask learning for lysine methylation site discovery"

Table S.2: Results of mass spectrometry analysis

| | Uniprot ID | Name | Position | Score | Unmethylated peptide detected | Methylated peptide detected | Hit |
|---|---|---|---|---|---|---|---|
| 1 | Q8N8Q3 | Endonuclease V | 170 | 0.905 | ✓ | ✓ | ✓ |
| 2 | Q9UBS4 | DnaJ homolog subfamily B member 11 | 66 | 0.931 | ✓ | ✓ | ✓ |
| 3 | P26641 | Elongation factor 1-gamma | 428 | 0.936 | ✓ | ✓ | ✓ |
| 4 | Q16836 | Hydroxyacyl-coenzyme A dehydrogenase, mitochondrial | 249 | 0.9 | ✘ | ✓ | ✓ |
| 5 | O14966 | Ras-related protein Rab-7L1 | 20 | 0.923 | ✘ | ✓ | ✓ |
| 6 | P58546 | Myotrophin | 11 | 0.916 | ✓ | ✘ | ✘ |
| 7 | Q9BSD7 | Cancer-related nucleoside-triphosphatase | 73 | 0.923 | ✘ | ✓ | ✓ |
| 8 | Q9NXW2 | DnaJ homolog subfamily B member 12 | 177 | 0.913 | ✘ | ✓ | ✓ |
| 9 | P61758 | Prefoldin subunit 3 | 59 | 0.971 | ✘ | ✘ | ? |
| 10 | Q9H7S9 | Zinc finger protein 703 | 141 | 0.897 | ✓ | ✘ | ✘ |
| 11 | Q7Z6K5 | Arpin | 203 | 0.958 | ✓ | ✓ | ✓ |
| 12 | P60228 | Eukaryotic translation initiation factor 3 subunit E | 407 | 0.906 | ✓ | ✓ | ✓ |
| 13 | P02794 | Ferritin heavy chain | 120 | 0.964 | ✓ | ✓ | ✓ |
| 14 | P56470 | Galectin-4 | 83 | 0.906 | ✘ | ✓ | ✓ |
| 15 | Q9NV56 | MRG/MORF4L-binding protein | 200 | 0.977 | ✘ | ✓ | ✓ |
| 16 | P27348 | 14-3-3 protein theta | 157 | 0.97 | ✓ | ✓ | ✓ |
| 17 | Q9BV29 | Coiled-coil domain-containing protein 32 | 98 | 0.913 | ✓ | ✓ | ✓ |
| 18 | P25787 | Proteasome subunit alpha type-2 | 50 | 0.957 | ✘ | ✘ | ? |
| 19 | Q9Y6G3 | Large ribosomal subunit protein mL42 | 114 | 0.935 | ✓ | ✓ | ✓ |
| 20 | P46734 | Dual specificity mitogen-activated protein kinase kinase 3 | 340 | 0.937 | ✓ | ✓ | ✓ |
| 21 | P25686 | DnaJ homolog subfamily B member 2 | 59 | 0.934 | ✘ | ✘ | ? |
| 22 | Q9NS73 | MAP3K12-binding inhibitory protein 1 | 208 | 0.965 | ✓ | ✓ | ✓ |
| 23 | Q9P0L0 | Vesicle-associated membrane protein-associated protein A | 24 | 0.917 | ✓ | ✓ | ✓ |
| 24 | O75608 | Acyl-protein thioesterase 1 | 105 | 0.913 | ✘ | ✓ | ✓ |
| 25 | O75934 | Pre-mRNA-splicing factor SPF27 | 218 | 0.932 | ✘ | ✓ | ✓ |
| 26 | P11233 | Ras-related protein Ral-A | 159 | 0.955 | ✓ | ✓ | ✓ |
| 27 | Q9NX63 | MICOS complex subunit MIC19 | 173 | 0.924 | ✘ | ✓ | ✓ |
| 28 | O15144 | Actin-related protein 2/3 complex subunit 2 | 256 | 0.909 | ✓ | ✓ | ✓ |
| 29 | Q96QD9 | UAP56-interacting factor | 261 | 0.976 | ✘ | ✘ | ? |

| | Uniprot ID | Name | Position | Score | Unmethylated peptide detected | Methylated peptide detected | Hit |
|---|---|---|---|---|---|---|---|
| 30 | P59998 | Actin-related protein 2/3 complex subunit 4 | 166 | 0.969 | ✘ | ✘ | ? |
| 31 | Q8NEF9 | Serum response factor-binding protein 1 | 378 | 0.942 | ✓ | ✓ | ✓ |
| 32 | O43505 | Beta-1,4-glucuronyltransferase 1 | 394 | 0.942 | ✘ | ✓ | ✓ |
| 33 | Q8IY31 | Intraflagellar transport protein 20 homolog | 34 | 0.923 | ✓ | ✓ | ✓ |
| 34 | Q9NWQ9 | Uncharacterized protein C14orf119 | 128 | 0.916 | ✓ | ✓ | ✓ |
| 35 | P26447 | Protein S100-A4 | 18 | 0.947 | ✓ | ✘ | ✘ |
| 36 | Q96KB5 | Lymphokine-activated killer T-cell-originated protein kinase | 8 | 0.984 | ✘ | ✘ | ? |
| 37 | P51809 | Vesicle-associated membrane protein 7 | 172 | 0.94 | ✓ | ✓ | ✓ |
| 38 | Q15286 | Ras-related protein Rab-35 | 189 | 0.969 | ✓ | ✓ | ✓ |
| 39 | P13073 | Cytochrome c oxidase subunit 4 isoform 1, mitochondrial | 78 | 0.907 | ✓ | ✓ | ✓ |
| 40 | P27216 | Annexin A13 | 284 | 0.963 | ✓ | ✓ | ✓ |
| 41 | Q8NA97 | Putative uncharacterized protein FER1L6-AS1 | 84 | 0.91 | ✘ | ✓ | ✓ |
| 42 | A0A024R1R8 | Translation machinery-associated protein 7B | 8 | 0.962 | ✘ | ✘ | ? |
| 43 | Q86SX6 | Glutaredoxin-related protein 5, mitochondrial | 151 | 0.93 | ✓ | ✓ | ✓ |
| 44 | Q5VT25 | Serine/threonine-protein kinase MRCK alpha | 1647 | 0.954 | ✓ | ✓ | ✓ |
| 45 | O75129 | Astrotactin-2 | 1337 | 0.917 | ✓ | ✓ | ✓ |
| 46 | Q13043 | Serine/threonine-protein kinase 4 | 480 | 0.964 | ✓ | ✓ | ✓ |
| 47 | Q13823 | Nucleolar GTP-binding protein 2 | 721 | 0.933 | ✘ | ✘ | ? |
| 48 | Q96IM9 | DPY30 domain-containing protein 2 | 158 | 0.925 | ✘ | ✓ | ✓ |
| 49 | Q8IX90 | Spindle and kinetochore-associated protein 3 | 389 | 0.954 | ✓ | ✓ | ✓ |
| 50 | O00300 | Tumor necrosis factor receptor superfamily member 11B | 255 | 0.912 | ✓ | ✓ | ✓ |
| 51 | P0DME0 | Protein SETSIP | 13 | 0.962 | ✘ | ✓ | ✓ |
| 52 | Q8NFR3 | Serine palmitoyltransferase small subunit B | 66 | 0.909 | ✓ | ✓ | ✓ |
| 53 | Q15506 | Sperm surface protein Sp17 | 49 | 0.897 | ✓ | ✓ | ✓ |
| 54 | Q9BVG4 | Protein PBDC1 | 63 | 0.939 | ✓ | ✓ | ✓ |
| 55 | Q0VDE8 | Adipogenin | 75 | 0.944 | ✘ | ✓ | ✓ |
| 56 | Q9UHD4 | Lipid transferase CIDEB | 213 | 0.946 | ✓ | ✓ | ✓ |
| 57 | A4D1E9 | GTP-binding protein 10 | 376 | 0.935 | ✓ | ✓ | ✓ |
| 58 | Q9NUV7 | Serine palmitoyltransferase 3 | 530 | 0.908 | ✓ | ✓ | ✓ |
| 59 | A2PYH4 | Probable ATP-dependent DNA helicase HFM1 | 1395 | 0.914 | ✘ | ✘ | ? |
| 60 | P20472 | Parvalbumin alpha | 37 | 0.975 | ✘ | ✓ | ✓ |
| 61 | Q8NE09 | Regulator of G-protein signaling 22 | 1254 | 0.942 | ✘ | ✓ | ✓ |

|  | Uniprot ID | Name | Position | Score | Unmethylated peptide detected | Methylated peptide detected | Hit |
|---|---|---|---|---|---|---|---|
| 62 | Q9UF47 | DnaJ homolog subfamily C member 5B | 184 | 0.933 | ✓ | ✓ | ✓ |
| 63 | P98175 | RNA-binding protein 10 | 915 | 0.942 | ✓ | ✓ | ✓ |
| 64 | Q5TC84 | Opioid growth factor receptor-like protein 1 | 344 | 0.918 | ✓ | ✓ | ✓ |
| 65 | Q92901 | Ribosomal protein uL3-like | 297 | 0.952 | ✘ | ✓ | ✓ |
| 66 | Q9HAV7 | GrpE protein homolog 1, mitochondrial | 138 | 0.923 | ✘ | ✓ | ✓ |
| 67 | P0DP57 | Secreted Ly-6/uPAR domain-containing protein 2 | 66 | 0.922 | ✓ | ✓ | ✓ |
| 68 | Q6P2I3 | Oxaloacetate tautomerase FAHD2B, mitochondrial | 18 | 0.929 | ✓ | ✓ | ✓ |
| 69 | B2CW77 | Killin | 75 | 0.957 | ✘ | ✓ | ✓ |
| 70 | O75223 | Gamma-glutamylcyclotransferase | 181 | 0.976 | ✓ | ✘ | ✘ |
| 71 | Q14914 | Prostaglandin reductase 1 | 318 | 0.955 | ✓ | ✓ | ✓ |
| 72 | P06276 | Cholinesterase | 586 | 0.934 | ✓ | ✓ | ✓ |
| 73 | Q6IQ20 | N-acyl-phosphatidylethanolamine-hydrolyzing phospholipase D | 18 | 0.906 | ✓ | ✓ | ✓ |
| 74 | O95372 | Acyl-protein thioesterase 2 | 69 | 0.918 | ✘ | ✓ | ✓ |
| 75 | Q6DD88 | Atlastin-3 | 399 | 0.958 | ✘ | ✘ | ? |
| 76 | Q9BZE2 | tRNA pseudouridine(38/39) synthase | 407 | 0.961 | ✘ | ✘ | ? |
| 77 | Q8N118 | Cytochrome P450 4X1 | 504 | 0.938 | ✘ | ✓ | ✓ |
| 78 | Q8WUD6 | Cholinephosphotransferase 1 | 383 | 0.926 | ✘ | ✓ | ✓ |
| 79 | Q96PM5 | RING finger and CHY zinc finger domain-containing protein 1 | 239 | 0.934 | ✓ | ✓ | ✓ |
| 80 | Q99757 | Thioredoxin, mitochondrial | 147 | 0.93 | ✘ | ✓ | ✓ |
| 81 | Q08623 | Pseudouridine-5′-phosphatase | 123 | 0.93 | ✓ | ✓ | ✓ |
| 82 | Q96CQ1 | Solute carrier family 25 member 36 | 174 | 0.924 | ✘ | ✓ | ✓ |
| 83 | Q8NGY0 | Olfactory receptor 10X1 | 325 | 0.975 | ✘ | ✘ | ? |
| 84 | Q9Y3V2 | RWD domain-containing protein 3 | 267 | 0.899 | ✓ | ✓ | ✓ |
| 85 | Q9ULV4 | Coronin-1C | 19 | 0.931 | ✘ | ✘ | ? |
| 86 | Q92499 | ATP-dependent RNA helicase DDX1 | 702 | 0.952 | ✓ | ✘ | ✘ |
| 87 | Q9H3Q1 | Cdc42 effector protein 4 | 114 | 0.973 | ✓ | ✘ | ✘ |
| 88 | Q9NSI8 | SAM domain-containing protein SAMSN-1 | 68 | 0.915 | ✘ | ✘ | ? |
| 89 | P61244 | Protein max | 24 | 0.956 | ✘ | ✘ | ? |
| 90 | Q8N4H5 | Mitochondrial import receptor subunit TOM5 homolog | 46 | 0.979 | ✘ | ✘ | ? |
| 91 | P25705 | ATP synthase F(1) complex subunit alpha, mitochondrial | 539 | 0.947 | ✘ | ✘ | ? |
| 92 | P28340 | DNA polymerase delta catalytic subunit | 1007 | 0.942 | ✘ | ✘ | ? |
| 93 | P49137 | MAP kinase-activated protein kinase 2 | 371 | 0.947 | ✘ | ✘ | ? |

| | Uniprot ID | Name | Position | Score | Unmethylated peptide detected | Methylated peptide detected | Hit |
|---|---|---|---|---|---|---|---|
| 94 | Q5MAI5 | Cyclin-dependent kinase-like 4 | 33 | 0.93 | ✘ | ✘ | ? |
| 95 | P30084 | Enoyl-CoA hydratase, mitochondrial | 101 | 0.972 | ✘ | ✘ | ? |
| 96 | Q9BTT4 | Mediator of RNA polymerase II transcription subunit 10 | 82 | 0.915 | ✘ | ✘ | ? |
| 97 | P08590 | Myosin light chain 3 | 107 | 0.967 | ✘ | ✘ | ? |
| 98 | P0C0P6 | Neuropeptide S | 80 | 0.987 | ✘ | ✘ | ? |
| 99 | Q96IG2 | F-box/LRR-repeat protein 20 | 408 | 0.897 | ✘ | ✘ | ? |
| 100 | Q32NC0 | UPF0711 protein C18orf21 | 85 | 0.96 | ✘ | ✘ | ? |
